# Temporally correlated active forces drive segregation and enhanced dynamics in chromosome polymers

**DOI:** 10.1101/2023.04.23.528410

**Authors:** Sumitabha Brahmachari, Tomer Markovich, Fred C. MacKintosh, José N. Onuchic

## Abstract

Understanding the mechanisms governing the structure and dynamics of flexible polymers like chromosomes, especially, the signatures of motor-driven active processes is of great interest in genome biology. We study chromosomes as a coarse-grained polymer model where microscopic motor activity is captured via an additive temporally persistent noise. The active steady state is characterized by two parameters: active force, controlling the persistent-noise amplitude, and correlation time, the decay time of active noise. We find that activity drives correlated motion over long distances and a regime of dynamic compaction into a globally collapsed entangled globule. Diminished topological constraints destabilize the entangled globule, and the active segments trapped in the globule move toward the periphery, resulting in an enriched active monomer density near the periphery. We also show that heterogeneous activity leads to the segregation of the highly dynamic species from the less dynamic one, suggesting a role of activity in chromosome compartmental segregation. Adding activity to experimental-data-derived structures, we find active loci may mechanically perturb and switch compartments established via epigenetics-driven passive self-association. The key distinguishing signatures of activity are enhanced apparent diffusivity, exploration of all the dynamic regimes (sub-diffusion, effective diffusion, and super-diffusion) at various lag times, and a broadened distribution of observables like the dynamic exponents.

## I. INTRODUCTION

Active processes, driven by molecular motors that consume energy (ATP) and exert persistent forces on biopolymers, are ubiquitous and crucial for sustaining cellular life. While motor-driven phenomena like network contractility [1–3] and motility [4–7] are well studied for stiff cytoskeletal polymer networks, the mechanical consequences of activity remain poorly explored for flexible polymers in the context of genome organization.

Folded chromosomes occupy mutually exclusive territories [8–10], where specific pairs of segments are more likely to be three-dimensional neighbors than others [11–15]. Chromatin bearing distinct epigenetic markers of transcriptional activity (euchromatin and heterochromatin) preferentially interact among themselves, forming A and B compartments [11, 16]. A characteristic structural feature is the organization of B compartments inside, and A compartments towards the periphery of chromosome territories [9, 17, 18]. Data-driven effective-equilibrium polymer models, via optimizing the inter-monomer-interaction energies, generate ensembles of folded structures consistent with experimental observations like contact frequencies [18–27]. These models, however, lack a direct connection to physical driving and are insufficient to explore the motor-driven aspects of chromosome structure and dynamics.

Chromosome dynamics is typically sub-diffusive at short time scales (seconds), as expected for a polymer [28–31]. However, there is significant heterogeneity in loci dynamics, reflected in altered mobility subgroups within the distribution of apparent diffusion constants [32–36]. In the effective-equilibrium approach, the altered mobility has been reasoned to arise from local confinement and/or the local compaction state [33, 37]. These structure-centric approaches are limited to only reducing the apparent diffusivity as compared to a free Brownian particle. The possibility of motor-activity-driven enhancement of dynamics is beyond such approaches. Noteworthy, some chromatin loci have been observed to move super-diffusively at intermediate time scales [35, 38, 39], which is at odds with any effective-equilibrium or passive approach. Active polymer models with motor-induced persistent forces have indeed shown the emergence of super-diffusive dynamics [40–44]. However, active control of chromatin structure is poorly understood.

We propose an active chromosome model where motor-generated forces within a coarse-grained locus contribute to its mobility. Within the active paradigm, coarsegrained motor activity contributes an additive noise that is temporally correlated (see Eq. (1) below). This contrasts active polymer systems constructed with different temperature particles, where the noise is temporally uncorrelated [45–49]. Active dynamics span sub-diffusion, effective diffusion, and super-diffusion. The apparent diffusion of a locus at long lag times is enhanced proportionally to the activity at the locus. Notably, activity not only introduces dynamical heterogeneity, but also affects the chromosome structure. Processes like transcription [50, 51], chromatin remodeling [52, 53], and loop extrusion [54, 55], that rely on ATP-consuming motors and exert forces on chromatin, are the hypothesized sources of activity.

A simplified model of a chromosome as a confined active polymer with soft self-avoidance shows that highly persistent or correlated motor activity typically expands the polymer and increases accumulation at the confinement boundaries. However, if the active kicks are not strong enough to overcome the entanglement constraints presented by chain connectivity and soft repulsion, the highly correlated active kicks lead to a collapsed globule state stabilized by the polymer entanglements. While the swelling and enrichment of polymer at the boundaries have been observed in active systems with different temperature particles [45, 46], the entangled globule is specific to long temporal correlations of the active noise, resembling the Motility-Induced Phase-Separated (MIPS) state observed in active colloids [56, 57]. Within the presented active model, these biologically relevant structural characteristics are dynamics-driven emergent properties, which we dissect further for mechanistic understanding. On the contrary, in effective-equilibrium models, these structures result from passive potentials like inter-monomer interactions [18, 19, 23–26, 58–60]. Interestingly, information on the steady-state structure alone cannot distinguish between active and passive models. The active steady-state polymer conformations may look identical to the passive ones of a completely unrelated system. Such as the collapsed globule of a purely self-avoiding active polymer may structurally disguise as a passive polymer with strong inter-monomer attractions. This highlights a potential ambiguity in deciphering the underlying mechanism from structural ensembles alone. However, signatures of enhanced dynamics, like super-diffusion or high apparent diffusivity, and correlated motion over long spatiotemporal scales unambiguously point towards an active system. While passive physical potentials have successfully recapitulated structures of chromosome compartments[18, 19, 23–26, 60], the presented model suggests that persistent motorized forces are a source of dynamic regulation of compartmentalization. We hope this work will lead to new experiments investigating the motor-driven dynamic aspects of chromosome compartments.

The layout of the article is as follows. We first describe the active polymer model (Fig. 1), then investigate the steady state of a confined, active homopolymer (Fig. 2). We explain the mechanisms of active regulation of the active structure and dynamics. To understand the consequences of heterogeneous activity, we simulate an active-passive block copolymer, uncovering features like phase segregation of active and passive monomers (Fig. 3).

**FIG. 1.**
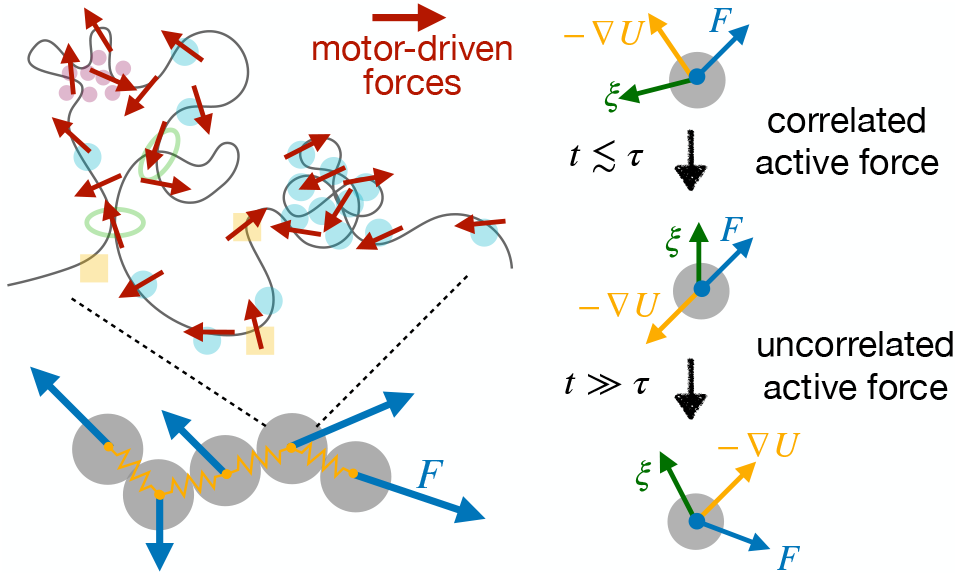
Schematic active polymer model. Microscopic motors exerting forces on the polymer are depicted as small red arrows. Each coarse-grained monomer containing distributed point forces experiences an active noise with amplitude *F*, such that the net active force on the polymer vanishes. At each time step, every monomer experiences three kinds of forces (Eq. (1)): temporally uncorrelated thermal noise *ξ*, passive forces derived from the polymer potential −∇*U*, and the active force *F*. The active kicks are correlated over a time scale *τ*, the correlation time. When averaged over long lag times (*t* ≫ *τ*), the correlation of the active kicks vanishes (Eq. (2))

**FIG. 2.**
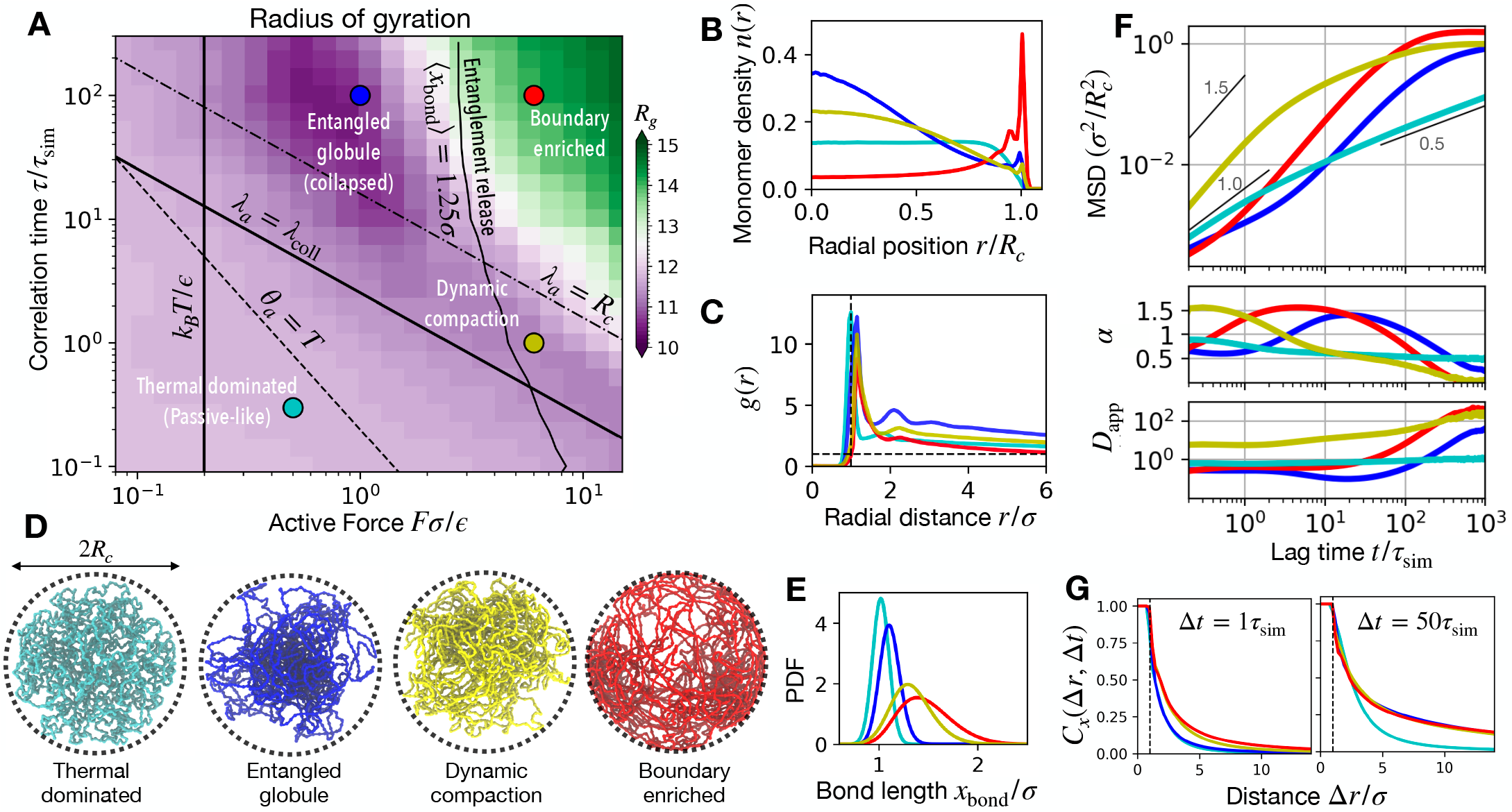
Self-avoiding active polymer in spherical confinement. (A) Active regime diagram spanned by the active force *F* (*ϵ/σ* units) and the correlation time *τ* (*τ*_sim_ units). The colors depict radius of gyration (*R*_*g*_) of the polymer (*σ* units). The active steady state (*θ*_*a*_ *> T*) shows compaction when the mean persistent path *λ*_*a*_ ≡ *Fτ/γ*, is longer than the mean collision distance between the monomers *λ*_coll_ = *σ/ϕ*^1*/*3^, where *ϕ* is the confinement volume fraction. This compaction is characterized by short-lived (dynamic) locally collapsed clusters. Long correlation times lead to a globally collapsed, entangled globule state. When *λ*_*a*_ is greater than the confinement dimension *R*_*c*_, the monomers have a tendency to get trapped at the boundary wall. However, release of entanglements is necessary to destabilize the competing entangled-globule state in order to enrich the boundaries. Hence only when the bonds are stretched by the active force (mean bond distance ⟨*x*_bond_⟩ *>* 1.25*σ*), leading to entanglement release, does the boundary-enriched state appear. Representative parameter sets for the four regimes are shown in colored circles for which various observables are plotted: thermal dominated (*F* = 0.5, *τ* = 0.3, cyan), entangled globule (*F* = 1, *τ* = 100, blue), dynamic compaction (*F* = 6, *τ* = 1, yellow), and boundary enriched (*F* = 6, *τ* = 100, red). We used *k*_*B*_ *T* = 0.2*ϵ* for all simulations. (B) Radial profile of the monomer number density *n*(*r*). (C) Radial distribution function *g(r)* shows a peak at 1*σ* corresponding to self-avoidance. Only the entangled globule state shows prominent successive peaks reflecting the collapsed state and an increased solid-like behavior. (D) Representative simulation snapshots where the dotted line represents the spherical confinement with radius *R*_*c*_. Note that the collapsed structures may appear as a passive polymer with self-adhesion, while the boundary-enriched state might resemble a passive polymer with boundary adhesion. This refers to the ambiguity in deciphering the underlying mechanism from structural ensembles alone. (E) Bond length distribution showing the increase in the mean bond length ⟨*x*_bond_⟩ for higher active force *F*. (F) Mean-squared displacement (MSD), normalized by the confinement dimensions, versus lag time *t* (*τ*_sim_ units), where the scaling exponents 0.5, 1.0, and 1.5 are drawn for comparison. The dynamic exponent *α*, and the apparent diffusivity *D*_app_, are plotted in the subpanels below. Active steady states span sub-diffusion, effective diffusion, and super-diffusion. The exponent *α* goes to zero at long lag times due to the confinement-induced saturation of MSD. (G) Correlation of monomer displacements measured over Δ*t*, plotted as a function of the distance between the monomers Δ*r*. Active steady states show enhanced correlated motion.

**FIG. 3.**
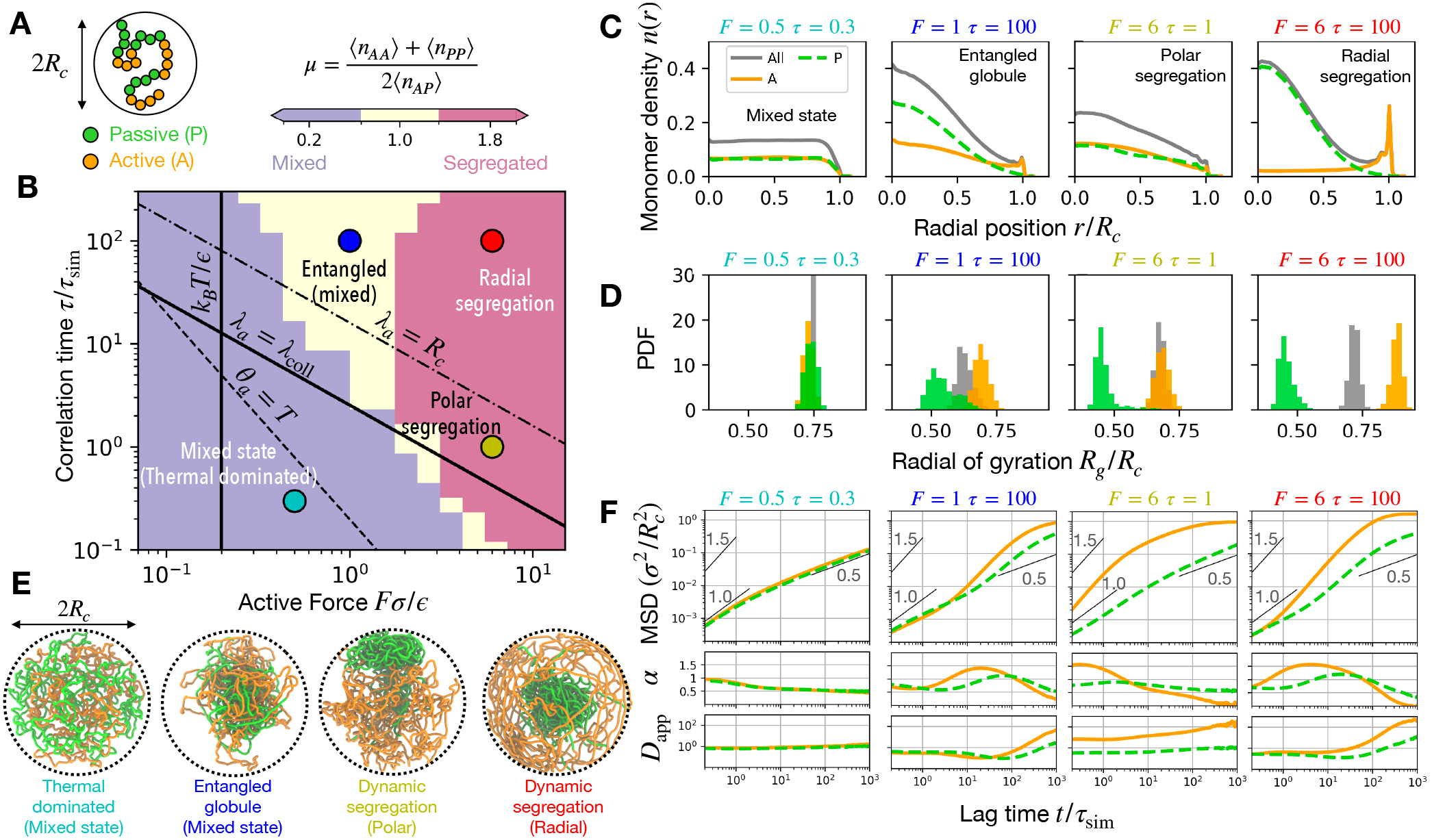
Phase separation of active and passive blocks of a confined self-avoiding polymer. (A) Schematic of the confined block copolymer with active (*A*) and passive (*P*) blocks with orange and green colors respectively. (B) Active regime diagram, where colors represent the phase separation coefficient *µ*, defined as the abundance of homotypic (*A* − *A* and *P* − *P*) over heterotypic (*A* − *P*) contacts. When *µ >* 1 (pink), there is segregation of the *A* and *P* blocks, while *µ <* 1 (purple) signifies a mixed state. The active steady state generally shows a tendency to segregate the active and passive blocks due their altered dynamics. However, the entangled globule state is only marginally segregated due to topological constraints. Given the active force is strong enough to release entanglements, activity with *λ*_*a*_ *> R*_*c*_ segregates the active and passive blocks radially. Otherwise, the passive blocks are segregated to a pole of the confinement. The four colored circles depict parameter sets for the four distinct regimes: thermal dominated (*F* = 0.5, *τ* = 0.3, cyan), entangled globule (*F* = 1, *τ* = 100, blue), polar segregation (*F* = 6, *τ* = 1, yellow), and radial segregation (*F* = 6, *τ* = 100, red). We used *k*_*B*_ *T* = 0.2*ϵ* for all simulations. (C) Radial profile of monomer density *n*(*r*) for active (orange, solid lines), passive (green dashed lines), and all monomers (grey solid lines). (D) Radius of gyration of the active (orange), passive (green), and all monomers (grey). (E) Representative simulation snapshots with *A* and *P* monomers colored in orange and green respectively. (F) Mean-squared displacement (MSD), the dynamic exponent (*α*), and apparent diffusivity (*D*_app_) versus lag time, for active and passive monomers corresponding to the four regimes. The dynamics of the active monomers are the same as described before (Fig. 2C). The passive monomers deviate from a purely passive behavior due to their polymer connectivity with the highly dynamic active monomers.

Building upon the intuition from the homopolymer and the block copolymer models, we discuss the model’s relevance to chromosomes. By adding active loci to a passive chromosome model (MiChroM [19, 61]), we explore active perturbation to the effective-equilibrium structures (Fig. 4). Finally, we conclude with an overview and discussion.

**FIG. 4.**
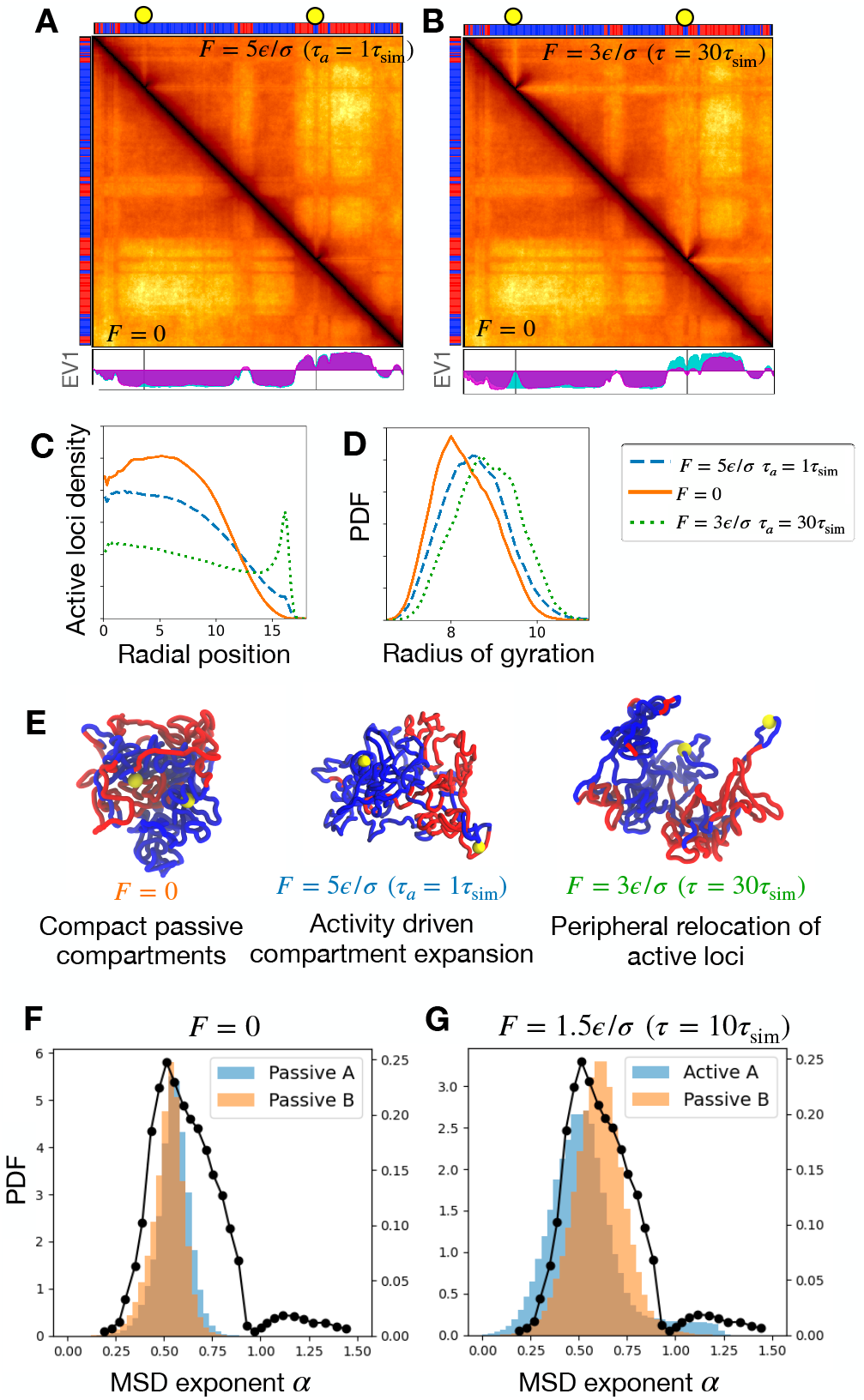
Activity expands passive compartments and contributes to heterogeneity in dynamics. (A) Simulated contact maps showing a 30 Mb segment (50 - 80 Mb) of the GM12878 chromosome 10. The lower triangle corresponds to the passive structure (*F* = 0), while the upper triangle corresponds to the active steady state with *F* = 5 and *τ* = 1. The thermal temperature is *k*_*B*_ *T* = 1.0*ϵ*. (B) Same as (A) with activity corresponding to a longer mean persistent time *F* = 3 and *τ* = 30. The bars on the top and left of the contact maps show the A (red) and B (blue) type beads that drive A/B compartments via passive phase separation. The yellow circles denote the two active sites in the segment. The principal eigenvectors (EV1) are shown in the subpanel below, where the passive (active) EV1 is colored in cyan (magenta). (C) Number density of active loci (including all the seven loci) plotted against radial position (*σ* units). (D) Radius of gyration (*σ* units) of the 30 Mb segment plotted in the contact maps above. (E) Simulation snapshots of the 30 Mb segment containing two active loci shown as yellow spheres. The A-type (B-type) regions are shown in red (blue). (F) The distribution of MSD exponents *α* for chromosome 10 simulated with passive MiChroM (*F* = 0) measured over lag times 1-100*τ*_sim_. (G) Same as (F) but with active *A* compartments (*F* = 1.5*ϵ/σ, τ* = 10*τ*_sim_). The black line with points (plotted on the right *y*-axis) corresponds to experiments in Human U2OS cells [35].

## II. ACTIVE POLYMER MODEL

The active polymer model hypothesizes the activity of molecular motors as point forces [41, 43, 44, 62–66]. As these are internally generated and not external body forces, they must be balanced, with no net force. We ensure this in our coarse-grained model with multiple active point forces that are randomly oriented (Fig. 1) and average to zero both on larger network scales and when averaged over long times. However, the motor binding kinetics makes the active force correlated over short time scales, set by the motor-residence time. This results in noise-like active kicks that are temporally correlated. The temporal decay of correlations is determined by the distribution of residence times of the motors. If motor (un)binding events are independent, the residence times follow a Poisson distribution giving an exponentially correlated noise. Following previous approaches [41, 43, 64, 66, 67], we assume that the active force correlation decays exponentially, characterized by a single time scale *τ*, the correlation time.

The overdamped equation of motion of the *n*-th active monomer in a thermal bath at temperature *T* reads as follows.

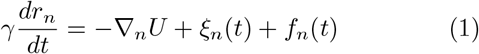

Here, *γ* is the drag coefficient, and −∇ _*n*_*U* represents the passive forces derived from the polymer potential, like the harmonic bonds between nearest neighbors. Random forces from thermal fluctuations, represented by *ξ*_*n*_, are uncorrelated: ⟨*ξ*_*n*_(*t*)*ξ*_*m*_(*t*^*′*^)⟩ = 2*γk*_*B*_*Tδ*(*t*−*t*^*′*^)*δ*_*mn*_, where *k*_*B*_ is the Boltzmann constant. The active noise *f*_*n*_, unlike thermal fluctuations, is correlated.

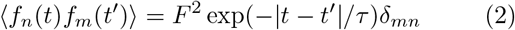

The amplitude of active noise *F*, a free parameter in the model, is expected to scale with the number of motors within the monomer. Note, we assume that the active forces of the neighboring coarse-grained monomers are uncorrelated. We ignore explicit correlations of the active force along the coarse-grained polymer contour, which may be of biological interest [48], and is left for future work. Including the effect of torque-inducing motors is yet another biologically relevant future possibility [68].

The technique of representing activity as a correlated noise has been discussed both in the context of motor activity in the cytoskeleton [62, 64, 67, 69, 70], as well as for active colloidal solutions, where the term Active Ornstein-Uhlenbeck Particles (AOUPs) is used [57, 71]. The steady state of AOUPs shows remarkable features such as inhomogeneous density profiles in confinement and Motility-Induced Phase Separation (MIPS): self-avoiding AOUPs segregate into a dense phase, mimicking effective attractive interactions [56]. The AOUP model has also been adopted to study Active Rouse polymers [41, 43, 72]. These polymer models focus on dynamics, while structural features, such as the analog of MIPS, remain largely unexplored.

### 1. Mean persistent path and the active steady state

The steady state of the active polymer is governed by a competition between the active (*F, τ*) and passive (*T*) parameters. A metric of activity is the emergent length: *λ*_*a*_ ≡ *Fτ/γ*, which we call the *mean persistent path* of an active monomer. This is the average distance an active monomer moves persistently before changing direction. In the presence of thermal fluctuations of strength *T*, the active features dominate only when the active temperature-like quantity *θ*_*a*_ = *Fλ*_*a*_ is dominant: *θ*_*a*_ *> T*. We generally refer to the parameter regime with *θ*_*a*_ *> T* as the active steady state and *θ*_*a*_ *< T* as the thermal-dominated passive-like steady state (Figs. 2A, 3B).

## III. RESULTS

### A. Self-avoiding active homopolymer in confinement

The physics underlying the active model (Eq. (1)) has been explored in the context of Rouse chains [41, 43, 72], which is in agreement with our model (see Appendix and Fig. A1). Here, we study the behavior of an active homopolymer with biologically relevant constraints: self-avoidance and confinement (see Appendix A for details of the potentials). Fig. 2 shows the results for a self-avoiding polymer with *N* = 2000 monomers that is confined within a sphere of radius *R*_*c*_ = 16*σ*, such that the volume fraction: *ϕ*≡ *Nσ*^3^*/*(2*R*_*c*_)^3^ ≈6%, is in the physiological regime for confined chromatin [73]. Since we simulate single chromosome polymer, the confinement effectively mimics the confines of the chromosome territory. Here, *σ* is the monomer diameter used as a unit of distance, the unit of energy is *ϵ*, and the unit of time is *τ*_*sim*_ (see Appendix for details).

#### 1. Activity drives dynamic compaction in self-avoiding polymers

A prominent signature of the active steady state is the compaction of a self-avoiding polymer into a denser globule with a lower radius of gyration than the passive case (Fig. 2A). The compact state emerges when the mean persistent path *λ*_*a*_ exceeds the average spacing between monomers, given by the concentration-dependent distance: *λ*_*coll*_ ≈ *σ/ϕ*^1/3^. In this regime (*λ*_*a*_ *> λ*_*coll*_), active monomers even after colliding with each other maintain their active noise direction, resulting in them getting trapped in a dynamically driven dense state. The compact state is characterized by dense clusters that coexist with less dense regions, where the correlation time dictates the stability of the compact clusters.

#### 2. Activity correlated over long times establishes a collapsed, entangled globule

Activity with a long correlation time traps the polymer into a globally collapsed globule (Fig. 2A). The collapsed globule is stabilized by topological entanglements, making the globule a long-lived state. Topological or polymer entanglements refer to the hindrance of the polymer segments to pass through each other due to the combined effect of the soft self-avoidance between monomers and the polymer connectivity. The polymer segments on the periphery of the globule may reorient their average active noise direction to escape the collapsed state. This leads to some segments coexisting in a less dense state with the entangled collapsed globule (Fig. 2D). Such segments, however, after momentarily exploring the less dense space outside the globule, reencounter the dense state, thus keeping the globule from dissociating. The compact structures driven by activity are liquid-like, however, the entangled globule exhibits an enhanced solid-like packing and mechanical integrity (Fig. 2C). The collapsed globule state is destabilized when the active force amplitude is smaller than the thermal fluctuations, *F < k*_*B*_*T/σ*, since thermal fluctuations destroy the correlated motion driving this state.

The collapsed globule is reminiscent of the segregated state in active colloids exhibiting MIPS [56, 57]. The collapsed state occurs at a lower density in active polymers (*ϕ*∼0.05) compared to AOUPs (*ϕ*∼0.5) [57, 74]. This stems from polymer entanglements stabilizing the dense phase. Enhancement of active signatures for polymers compared to colloids with the same activity has been observed in simulations [46]. Semiflexible polymers without self-avoidance show activity-driven compaction because activity enhances transverse fluctuations that lead to a lower end-to-end distance [75]. However, this compaction mechanism differs from the self-avoidance and polymer connectivity-driven collapse into the entangled globule.

#### 3. Large active noise amplitudes destabilize the collapsed globule via stretching bonds and reducing entanglements

The collapsed, entangled globule gradually disappears as the active noise amplitude *F* is increased (Fig. 2A). Increasing the active force *F* increases the fraction of the polymer that coexists in the less dense space outside the globule, eventually leading to the complete destabilization of the dense state. This destabilization is due to the stretching of the bonds of the polymer by strong active kicks, which diminishes the topological constraints. Two non-neighboring segments with stretched bonds are more likely to pass through each other than get trapped in a dense state. We find that ≈25% increase in the average bond length is enough to destabilize the collapsed state (Fig. 2A, E).

Following our rationale, increasing the bond stiffness, which decreases the bond length and reinforces the entanglement constraints, stabilizes the collapsed state for higher active noise amplitudes (Figs. A4). The stability of the entangled globule state for low active noise amplitudes is governed by the thermal temperature. When *F < k*_*B*_*T/σ*, uncorrelated thermal noise dominates the correlated active noise, leading to destabilization of the collapsed, entangled state (Figs. 2A, A5). Note, although, the thermal noise domination makes the structure appear equilibrium-like, the dynamics at long correlation times can still show signatures of activity.

#### 4. Competition between mean persistent path and confinement dimension leads to boundary enriched polymer

A boundary-enriched state emerges for high active force and high correlation times, where the monomer density peaks at the boundary, is depleted at the center, and the polymer is expanded such that the radius of gyration assumes the size of the box (Fig. 2A, B). When the mean persistent path is longer than the confinement dimension, i.e., *λ*_*a*_ *> R*_*c*_, the active monomers collapse on the boundary wall. The active monomers with long correlation times keep pushing against the wall, while upon reorientation of their active force direction, they move in a correlated fashion until encountering the other side of the boundary. However, this state only appears when the entanglement-driven constraints are sufficiently weaker so that the polymer does not get trapped in the entangled globule state. Hence, the two criteria for the appearance of this state are the high force, necessary for the release of entanglements, and a high enough mean persistent path, necessary for getting trapped at the boundary (Fig. 2A). Boundary enrichment comes about from a competition between monomer activity and confinement, hence, self-avoidance and polymer connectivity are not necessary for establishing this state. As a result, active Rouse chains and active gas or colloidal particles particles (AOUPs) both exhibit emergence of boundary enrichment with appropriate mean persistent path (Figs. A2 and A3).

#### 5. Structural ensembles are insufficient to distinguish between effective-equilibrium and actively driven phenomena

The conformations of the active steady state may appear to belong to a completely unrelated passive system. For example, the collapsed state might look like a polymer with self-adhesive interactions (Fig. 2D). Or, the boundary-enriched structures may appear as though the monomers have attractive interactions with the confinement wall (Fig. 2D). This clearly suggests that structural ensembles are not enough to distinguish between the underlying active and passive mechanisms. However, the dynamics may hold key distinguishing features.

#### 6. Regimes of polymer dynamics

The dynamics in viscoelastic media is often characterized by anomalous diffusion, with the mean-squared displacement (MSD) given by ⟨(*r*_*n*_(*s* + *t*) −*r*_*n*_(*s*))^2^⟩ = *Dt*^*α*^, where the dynamical exponent of the lag time *t* classifies distinct dynamic regimes: diffusion for *α* = 1, subdiffusion for *α <* 1 and super-diffusion for *α >* 1. The coefficient *D* is an apparent diffusivity parameter. A free passive particle executes diffusion, where *D* is the diffusion constant proportional to the thermal temperature: *D*_free_ = *k*_*B*_*T/γ*.

Polymer relaxation spans multiple time scales that arise because fluctuations with longer wavelengths decay slower. The shortest wavelength fluctuations, corresponding to bond fluctuations, relax at short time scales *τ*_bond_∼*γ/k*, where *k* is the bond stiffness. On the other end of the spectrum, *τ*_Rse_ ≈ *N* ^2^*γ/*(*kπ*^2^) corresponds to the relaxation time of the longest wavelength [76] (see Appendix B; for Figs. 2 and 3, *τ*_bond_ ≈0.1*τ*_sim_ and *τ*_Rse_ ≈ 10^4^*τ*_sim_). The third important time scale is the active correlation time *τ*. An interplay of these time scales determines the dynamic signatures.

For lag times shorter than the bond relaxation time (*t* ≪ *τ*_bond_ *< τ*), passive polymer dynamics resembles free-particle diffusion (Fig. 2F). For lag times larger than the bond relaxation but smaller than the longest polymer relaxation time (*τ*_bond_ *< t < τ*_Rse_), passive dynamics exhibits sub-diffusion (*α <* 1) (Fig. 2F) [76]. This originates from each monomer having to drag a portion of the chain with it. Within this regime, active polymers may have substantially different signatures (Figs. 2F, 3F, A1). At lag times longer than *τ*_Rse_, passive motion returns to diffusion, corresponding to the polymer center-of-mass motion [76]. For chromosomes, *τ*_Rse_ is typically very long and experimentally inaccessible. For this reason, we have not included this regime in Figs. 2F and 3F. In the case of confined polymers, this may not be observed because MSD at such long lag times may be saturated due to the confinement. The dynamic regimes of the active self-avoiding polymer are similar to that of the active Rouse chain (see Appendix and Fig. A1) [41, 43], since all the dynamic signatures originate from a competition between the monomer activity and the nearest neighbor bonds in the polymer potential.

#### 7. Super-diffusion in active polymers

The leading order active contribution to MSD scales as quadratic in time ∼(*t/τ*)^2^, driving a super-diffusive regime [41, 43] (Appendix B). Whereas, at the short time scales (*t*≪ *τ*_bond_), the thermal contribution is linear in lag time. This results in the MSD asymptotically approaching the diffusive (linear) regime near zero lag times irrespective of activity (Fig. 2F).

The activity-driven super diffusion in the MSD curves appears when the ∼*t*^2^ active contribution exceeds the passive forces. The dominance of the active forces leads to a super-diffusive exponent when the lag time is comparable to the correlation time (Fig. 2F). For short correlation times (*τ*≈*τ*_bond_), the active polymer shows super-diffusion at the time scales of free-monomer dynamics (Fig. 2F).

For correlation times that are longer than the bond relaxation time (*τ*_bond_ *< τ < τ*_Rse_), the dynamics show a sub-diffusive regime at intermediate lag times: *τ*_bond_ *< t < τ* (Fig. 2F). This sub-diffusion is a result of compensation of the persistent active drive by the polymer potential [41]. Interestingly, the entangled globule shows an extended sub-diffusive regime due to the suppression of dynamics in a crowded environment (Fig. 2F).

Active polymer dynamics restores to sub-diffusion at lag times longer than the correlation time (Fig. 2F). Within our model, at these long lag times, the active noise becomes indistinguishable in its time dependence from thermal noise. However, the motion is enhanced in amplitude due to the active noise contribution, reflected in an increased apparent diffusivity.

#### 8. Activity typically enhances the apparent diffusivity

We compute an apparent diffusivity from the simulated MSD curves: *D*_app_ ≡ MSD*/t*^α^, where the exponent *α* ≡ *∂*(ln MSD)*/∂*(ln *t*). Note that the apparent diffusivity is a lag-time dependent coefficient that is equal to the diffusion constant when *α* = 1. For lag times shorter than the bond relaxation time *τ*_bond_, the apparent diffusivity for a passive polymer is set by the free-particle diffusion constant: *D*_app_ ≈ *k*_*B*_*T/γ* (Fig. 2F). While at lag times longer than the polymer relaxation time *t > τ*_Rse_ *> τ*, the passive apparent diffusivity is again a constant: *D*_app_≈ *k*_*B*_*T/*(*Nγ*), and is lower due to the center-of-mass motion [76].

The apparent diffusivity for active polymers shows enhancement over the corresponding passive behavior only for lag times longer than the correlation time (Fig. 2F). Interestingly, the active apparent diffusivity may also show suppression, such as in the entangled globule state. Competition between the active noise and the passive polymer potential underlies this suppression.

While super-diffusion is a unique characteristic of the active steady-state, investigating the dynamics at the appropriate lag time is essential to observe this. When lag times are longer than the active correlation time, the dynamics appear passive with a higher effective temperature (Fig. 2F). While lag times smaller than the correlation time may show suppressed dynamics akin to lower effective temperatures. Hence, observations of passive-like dynamic exponents do not rule out active forces, the apparent diffusivity and MSD exponents should be scrutinized at various lag times to investigate an active component.

Active dynamics bear another key signature: correlated motion of the monomers over long distances. Activity-driven compact clusters exhibit correlated motion, reflected in a shallower decay of the two-point displacement correlation function for the active steady states (Fig. 2G).

### B. Phase segregation of active and passive blocks of a confined self-avoiding polymer

To explore the effect of heterogeneity in activity along a polymer, we study a block copolymer with alternating active (*A*) and passive (*P*) blocks (Fig. 3A). The polymer has *N* = 2000 monomers and each alternating active and passive block of size *n* = 200 monomers. We use the same self-avoidance and confinement conditions (*ϕ* = 0.06) as before (see Appendix A). In the passive-like regime (*T > θ*_*a*_), the *A* and *P* blocks mix due to entropy and the dynamics is equilibrium-like (Fig. 3F). The active steady states (*T < θ*_*a*_) tend to show phase segregation of *A* and *P* blocks. Spatial separation of two species with different dynamics underlies this segregation. Highly dynamic active monomers occupy a larger share of the confinement volume, relegating the passive monomers into a compact segregated state. Interestingly, the spatial organization of the segregated phases is governed by active correlation time (Fig. 3B, C, E).

Activity-driven phase separation between the active and passive blocks is observed only when the active force is large enough to release entanglements (Fig. 3B). Phase separation is measured via the coefficient *µ*, defined as the ratio of homotypic to heterotypic pairwise contacts between inter-block monomers. A pairwise contact is defined when the inter-monomer distance is less than 1.5*σ* (Appendix A). When *µ >* 1 the active and passive blocks segregate, whereas *µ <* 1 denotes the mixed state.

The entangled globule state shows an overall collapse of the entire polymer, where the active and passive blocks are not well segregated (Fig. 3B-E). The regime corresponding to boundary enrichment of the active blocks (high *F* and high *τ*), shows strong phase separation (Fig. 3B, C). In this regime, the passive blocks are highly compacted and reside at the center, while the active blocks spread over the boundary wall (Fig. 3E). Phase separation is also observed in the regime of low mean persistent path (high *F* and low *τ*). This state is characterized by compact passive domains and expanded, highly dynamic active blocks that are segregated without any radial preference, in a polar fashion (Fig. 3B-E). Increasing the thermal temperature releases entanglements for passive blocks and leads to a boundary enrichment of passive monomers (Fig. A9).

The dynamics of the active monomers in the block copolymer basically resemble the regimes explored in the active homopolymer case (Figs. 2F and 3F). Interestingly, the dynamics of the passive blocks show deviation from passive behavior (Fig. 3F). The bonded interactions with the active monomers make the passive blocks more dynamic.

Phase separation of passive block copolymers is typically modeled using the Flory-Huggins approach, where blocks are assigned self and mutual interactions. The active phase separation presented here is fundamentally different, as it is dynamically driven. Mutually repulsive active monomers segregate from the passive monomers creating regions of high and low dynamics. Although the structural ensembles of a Flory-Huggins polymer may appear similar to the active phase segregated system, the dynamics are more heterogeneous and span multiple regimes for the activity-driven structures. The dynamic segregation within an active-passive mixture has been observed in other polymer models [47, 77].

### C. Implications for chromosomes

Chromosomes are active matter, where motorized processes like transcription, loop extrusion, and chromatin remodeling are continually driving the structure and dynamics. Our understanding of chromosome folding and the underlying mechanics has consolidated thanks to the emulsification of experimental data like Hi-C contact maps [11–13] and polymer models [18–27]. However, the existing chromosome models are predominantly effective-equilibrium, lacking direct signatures of activity like superdiffusive chromatin loci [35, 38, 39].

Here we use the presented active model to study chromosomes. The amplitude of the active noise is set by the typical stretching elasticity of chromatin: *F*≈1 −10 pN [78, 79]. The active correlation time *τ* is a measure of the motor-residence time. While chromatin remodellers typically show fast, sub-second dynamics [53], persistent bursts of gene activity, corresponding to RNA polymerases moving processively, may last for many minutes [80]. Loop extruding enzymes (SMC complexes) typically extrude a 50 kb DNA loop in tens of seconds [55]. Note, extrusion activity of loops shorter than a monomer size (≈ 50 kb) may be considered within this approach, whereas large loops spanning multiple monomers should be incorporated explicitly as large force dipoles. Hence, we argue that the physiological range for the correlation time is *τ*≈1 − 10^2^ seconds, where the higher (lower) end of the spectrum is associated with transcription (chromatin remodeling and extrusion of short loops). Consequently, at experimentally realizable time scales (1-100 seconds), we expect chromatin regions housing highly transcribed genes to show super-diffusive signatures, whereas, loci containing loop extruders or chromatin remodellers are expected to show sub-diffusion with an enhanced apparent diffusivity.

The eukaryotic genome is typically confined within the nucleus with a volume fraction ≈1 − 10% [73]. The hierarchical organization of the genome may introduce effective confinement with a similar volume fraction but at a smaller dimension. Such as the chromosome territories, which are about an order of magnitude smaller than the nuclear dimension, may effectively confine chromosomal loci, like genes or centromeres [10]. Hence there are multiple confinement lengthscales that may compete with the mean persistent path of an active chromosome locus.

#### 1. Activity-driven phase separation as a mechanism of chromosome compartmentalization

Compartmentalization is an important feature of chromosome folding in the interphase chromosomes of eukarya [11, 13, 16]. Compartments are globules formed by colocalization of sequentially distant genomic elements, reflected in the off-diagonal plaid-like patterns of the Hi-C maps. Hi-C-defined (sub)compartments are structural classes obtained via dimensionality reduction, e.g., Principal Component Analysis (PCA) of the pair-wise genomic-interaction patterns [11, 13]. These compartments are correlated to the cellular gene expression profile and show tissue- or cell-type-specific variability [12, 81–83]. However, the mechanics governing the relationship between structurally annotated compartments and gene expression is not understood.

The active model suggests phase-segregated structures, akin to A/B compartmentalization may arise from altered activity between A and B segments. The distinguishing characteristics of active compartments are their enhanced dynamics, elevated apparent diffusivities, and the possibility of a super-diffusive dynamic regime (Figs. 2C-E and 3F). Polymer-entanglement-based constraints stabilize these motor-driven compartments. Hence, increased activity of enzymes like type-II DNA topoisomerase, releasing topological constraints, is expected to destabilize these compartments.

A passive mechanism for compartmentalization, that has been widely explored, is short-range attractive interactions originating from the chromatin loci’s chemical nature (epigenetics) [18–27, 60]. Globules established via motor activity may act as a nucleation site for the three-dimensional spreading of epigenetic marks [84, 85]. This will lead to robust compartments that persist even when the motor activity ceases. Alternately, compartments established via passive attraction may be disrupted or perturbed by altered activity.

To investigate how passive compartments may respond to activity, such as increased transcription, we simulate active loci within the passive MiChroM model [19, 61]. Briefly, the optimized MiChroM potential contains favorable inter-monomer interactions that may be divided into two categories: first, the phase separation term that drives compartmentalization, and second, the ideal chromosome term that drives lengthwise compaction [19, 58]. We simulate chromosome 10 of Human GM12878 cell at 50 kb resolution (*N* = 2712) using Eq. (1), where the passive forces (−∇*U*) originate from the MiChroM potential (see Appendix C and Fig. A10). Additionally, we introduce seven active loci (five in B compartments and two in A compartments) that model increased motor activity at those sites (Appendix C). We then studied how the structural ensembles change when the active loci correspond to three distinct regimes: passive (*F* = 0); active with short mean persistent path (*F* = 5, *τ* = 1); and active with long mean persistent path (*F* = 3, *τ* = 30) (Fig. 4).

#### 2. Localized activity may expand compact passive compartments, induce compartment switching, and displace the active loci to chromosome periphery

Active loci exhibit enhanced dynamics, and the resulting agitation perturbs the compact compartments harboring the active loci. Active agitation opened up the compartments leading to a higher radius of gyration (Fig. 4D). The simulated Hi-C maps show local loss of contacts (Fig. 4A-B). Structural perturbation via active loci also reflects in a change in the principal eigenvector of the correlation matrix (EV1) (Fig. 4A-B).

Active loci with long mean persistent paths extend out of the compartment and tend to move towards the chromosome periphery (Figs. 4C, D, and A11). This leads to a light stripe in the simulated contact map, as the peripherally located active loci have limited interaction with the rest of the chromosome (Fig. 4B). Highly correlated activity may diminish the passive-compartment strength or altogether flip the compartment signature (EV1) at the active site (Fig. 4A, B). The Hi-C signature resembles the “jet”-like protrusions observed in experiments at localized loading sites of loop extruders (SMC complexes) [86].

Activity with a long mean persistent time may arise from persistent bursts of transcription lasting for many seconds to minutes. Positioning of transcriptionally active segments to the periphery of chromosome territories is experimentally well-documented [15, 59, 87–90]. Additionally, high transcriptional activity is correlated with long genes looping out of the chromosome territories [59]. The peripheral positioning of highly expressed genes leads to loss of contact with the rest of the chromosome, which has also been observed recently [91]. Activating transcription within a repressive compartment may drive compartment switching, which may ultimately lead to cell-fate transitions [12, 82]. The active model argues that the motorized mechanics of the transcription process directly contribute to all these phenomena. Modeling motor activity as higher temperature also shows the structural relocation of active monomers to the periphery [45, 92], which may be compared with the low correlation time activity (Fig.4A, C). However, the effect is stronger for activity with higher correlation times (Fig.4B, C).

#### 3. Activity increases the heterogeneity in chromatin dynamics

Passive MiChroM is useful for reconstructing the structure from Hi-C maps but the heterogeneity in chromatin dynamics is somewhat underrepresented (Fig. 4F). In fact, any passive model is incapable of recapitulating super-diffusive chromatin loci, represented by *α >* 1 in experimental data (Fig. 4F, G) [35, 39]. Adding activity to the A-compartments of passive MiChroM led to a substantial widening of the MSD exponent distribution, better recapitulating the experimental observations (Fig. 4G). Active simulations also exhibit the super-diffusive fraction, which is a feature of the active A compartments. Our model suggests that motor-driven active forces are an essential component in modeling the dynamics of chromosomes. While an important aspect of chromosome dynamics is derived just from its polymer nature, motor activity is the additional ingredient necessary to fully capture chromosome dynamics.

## IV. DISCUSSION

We have studied a coarse-grained polymer model of chromosomes that, in addition to the uncorrelated thermal noise, experiences a temporally correlated active noise (Eqs. (1) and (2)). This active noise models the effect of microscopic motors exerting forces inside the coarse-grained monomer (Fig. 1). Being internal, these forces must be balanced, with no net forces on the center of mass. At the molecular scale, one can think of such activity as point-like dipolar forces. We only consider network degrees of freedom in our model and not fluid stress propagation. In such an over-damped, Brownian dynamics model, forces must balance within the network. This balance, however, may only occur at scales of the order of the distance between entanglements or points of contact between non-neighboring segments in the network. Moreover, at such entanglement points, forces can branch, resulting in more complex multi-polar forces on coarse-grained scales, although with still vanishing net force on the center of mass. Thus, although the presented model allows for unbalanced active force at each monomer, the model does ensure that the net force on larger network scales is balanced, due to the large number of randomly oriented active force monopoles. Hence, there is no net momentum of the polymer center of mass in our model. We additionally verified the momentum conservation of the polymer network by implementing a variation of the model where the active noise of each monomer is modified such that the polymer center of mass is suppressed (Figs. A6, A7, A8). All the qualities of our results remain unchanged except for an enhanced suppression of the apparent diffusion constant of the center of mass of the active polymer in certain regimes.

We have presented a simplified characterization of the steady state of the active polymer using two parameters: the force *F* depicting the active noise amplitude and the correlation time *τ* controlling the temporal persistence of the active noise (Figs. 2A and 3B). The model exhibits diverse structural and dynamic properties. Active-polymer dynamics span sub-diffusion, effective diffusion, and super-diffusion, where the correlated active kicks underlie the super-diffusive regime (Figs. 2F and 3F). Importantly, non-equilibrium activity can result not only in enhanced monomer dynamics (e.g., super-diffusion) but also in structure formation that arises from stochastic active forces. Contrasting features like polymer swelling/collapse and radial phase segregation with higher monomer density towards the center/periphery, are some of the actively controlled structural aspects (Figs. 2B-D and 3C-E). The active steady-state structures may look similar to that of a passive system with attractive or repulsive interactions between monomers, however, the underlying mechanisms are completely different (Figs. 2D and 3E) the dynamics is essential to distinguish between active and passive mechanisms.

We propose motor-driven collapse as a mechanism of chromosome compartmentalization. These compartments are steric-hindrance and entanglement driven, such that they can be destabilized via enzymes like type-II DNA topoisomerases that release entanglements. Upon entanglement release, the segments with correlated activity, such as highly transcribed genes, move toward the periphery of chromosomes. This posits that the motorized mechanics of transcription may be contributing to the relocation of the highly transcribed genes to chromosome territory edges. We also found that phase-separated compartments established by passive attractive forces are destabilized by activity due to the enhanced local dynamics (Fig. 4). Recent modeling has proposed that activity correlated along the polymer chain may drive compartmentalization [48]. Our model shows that compartment-like compaction can also arise from an interplay of entanglements and temporally correlated activity. As mentioned before, the distinction between the active and passive mechanisms can be made based on measurements of dynamics [48].

Cell-type-specific variations in the genome architecture are reflected in the compartment structure [12, 81–83]. These observations posit a conundrum: do compartments drive gene expression or vice-versa? While it is conceivable that the distinct chemical microenvironments within compartments help recruit transcription machinery thus aiding gene expression, our results suggest there is a possible mechanical feedback wherein a highly transcribed locus may alter the compartment structure surrounding the locus. Investigating the causality between compartments and gene expression is key to deciphering the genome structure-to-function relationship.

Chromatin loci typically show sub-diffusion, which may be either passive or active in nature [28–31, 93, 94]. Interestingly, experiments have confirmed the presence of a wide distribution of dynamic exponents, including super-diffusion [35, 38, 39], which must be active in nature. We found that the passive chromosome model better recapitulated dynamic exponents when motor activity was added to the A compartments (Fig. 4G), suggesting motor activity as a driver of heterogeneity in dynamics. However, activity perturbed the structure to worsen agreement with Hi-C maps. This necessitates that the passive interactions be recalibrated in the presence of dynamics-satisfying motor activity, which is left for future work.

Our work is motivated in part by a large body of prior work on nonequilibrium aspects of cytoskeletal networks driven by molecular motor activity[1–3, 62, 64, 67, 70, 95]. Persistent active noise has been argued to effectively capture stochastic binding and force generation by microscopic motors in polymer networks, giving rise to organization, mechanics, and dynamics on various time scales [64, 67]. Models based on explicit motor-induced correlated forces upon stochastic binding have also been shown to recapitulate network contractility [3]. The present work builds on the prior approaches without explicit motor binding dynamics, where the motor binding/unbinding time scale is effectively captured by the active correlation time.

More recently, attention has turned to similar active matter aspects in chromosomes [40–43, 45, 46, 48, 49, 66, 96]. One class of active polymers corresponds to models with multiple temperatures. In this approach, monomers experience instantaneous thermal kicks, but there are at least two different types of monomers based on their effective temperature, i.e., “hot” and “cold” particles [45– 47, 77, 92, 97]. These models show structural features like phase segregation, polymer enrichment at confinement boundaries, and dynamics aspects like a broad apparent diffusivity. However, the possibility of a superdiffusive dynamic regime is beyond such an approach.

Active polymer models that explicitly model active force dipoles constitute the second class of active models [49, 98]. These models, utilizing instantaneous force dipoles, have shown a paradigm of structure and dynamics regulation by the dipolar strength. However, the dynamics lack any super-diffusive component as well. Important contributions have been made by studies incorporating hydrodynamic aspects along with dipolar activity that argue active stresses originating from the dipoles as the driver of convective flows in the chromatin [96, 99].

The treatment of activity as a temporally persistent noise, as done in the present work, has been explored in other models of flexible, Rouse-like polymers [41, 43, 72]. These models explain the origin of a super-diffusive dynamics component due to the active correlation time, where a longer correlation time, resembling longer motor residence times, drives a more prominent super-diffusive regime. This agrees with the explicit modeling of stochastic binding and force generation between motors and polymers, which shows an enhanced superdiffusive regime with higher motor concentrations and longer motor residence times [42]. The present work, however, goes beyond dynamics and has identified for the first time an entangled globule organization reminiscent of MIPS due to the combination of activity and repulsive interactions missing in prior Rouse-based models. One prior work [100] has studied self-avoidance and correlated active noise together in polymers but does not identify the novel structural and dynamic phases emerging in various activity regimes presented here, including the entangled globule phase.

Through mechanistic explanations of various active, steady states, the present work puts into perspective the observations of previous works like multiple-temperature activity as a special case of high force and low correlation time and thus widens the scope of possible structure and dynamic features of polymers driven by persistent activity. We hope this work will lead to many new non-equilibrium models of chromosomes, and aid in the design of future experiments investigating the signatures of motor activity in chromosomes.

## V. ACKNOWLEDGMENTS

This work was supported by the Center for Theoretical Biological Physics (CTBP) sponsored by the National Science Foundation (NSF Grants PHY-2019745, PHY-2210291 and DMR-2224030) and by the Welch Foundation (Grant C-1792). We are grateful for the computational resources provided by Advanced Micro Devices (AMD). JNO is a Cancer Prevention and Research Institute of Texas (CPRIT) Scholar in Cancer Research. TM acknowledges funding from the Israel Science Foundation (Grant No. 1356/22). SB acknowledges the Robert A. Welch Postdoctoral Fellow program. We would like to thank Peter Wolynes, Vinicius G. Contessoto, Antonio Oliveira Jr., and other members of CTBP for helpful discussions.

## Appendix A: Methods

### 1. Equation of motion

The overdamped equation of motion for the *n*-th monomer is given by Eq. (1). The active force *f*_*n*_(*t*) denotes the temporally correlated active noise. The temporal evolution of the active force for the *n*-th monomer is given as follows.

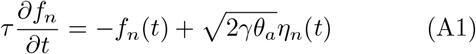

Here, *τ* is the correlation time of the active force, and *θ*_*a*_ ≡ *F* ^2^*τ/γ* is the temperature-like quantity associated with activity. The amplitude of the active noise is denoted by *F*. Finally, *η*_*n*_(*t*) is a delta-correlated stationary Gaussian process with zero mean: ⟨*η*(*t*)⟩ = 0, and ⟨*η*_*n*_(*t*)*η*_*m*_(*t*^*′*^)⟩ = *δ*(*t* − *t*^*′*^)*δ*_*mn*_.

Solving the equation of motion of the active force (Eq. (A1)), we get the following autocorrelation.

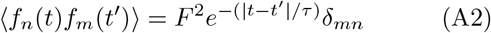

The active force between different monomers are always uncorrelated. At short lag times (*t* ≲ *τ*) the motor-induced noise for a monomer is correlated, leading to persistent dynamics. While at longer lag times (*t*≫ *τ*), the correlation vanishes exponentially.

### 2. Brownian Dynamics Simulations

We use Brownian Dynamics simulations where the positions are updated at each time step based on forces and positions of the previous time step. We integrate the overdamped Langevin equation of motion of a monomer (Eq. (1)) using the following two steps:

1. *Compute the net force*. Active forces at time *t* are computed from Eq. (A1):

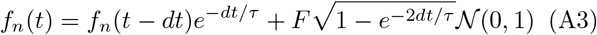

where 𝒩 (0, 1) is a random variable drawn from a Normal distribution with zero mean and unit standard deviation. Note, the active force does not depend on particle positions. The net force *h*_*n*_(*t*) also has a passive contribution. The passive force is computed from the interaction potential using monomer positions {*x*(*t*)}:

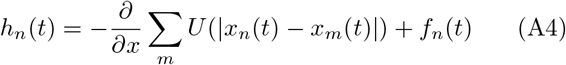
2. *Update positions*. Then, the monomer positions at time *t* + *dt* are obtained according to Eq. (1):

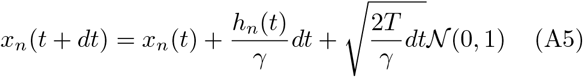

We simulate the equation of motion for each particle (Eq. (1)) using a custom integrator within the OpenMiChroM simulation package [61], which is built on top of OpenMM [101].

### 3. Simulation and physical units

The unit of distance is set by the monomer diameter *σ*. The simulation unit of energy is *ϵ*, such that forces are measured in *ϵ/σ*. The simulation unit of time is *τ*_*sim*_. We use a drag coefficient of *γ* = 1.0 *ϵτ*_*sim*_*/σ*^2^.

To calibrate with physical units, we use *ϵ* = *k*_*B*_*T*. The monomer diameter corresponding to about 50 kb chromatin is *σ* ≈ 100 nm. The apparent diffusion constant can be written as *D*_*app*_ = *k*_*B*_*T/γ* = *σ*^2^*/τ*_*sim*_. This equated with the experimentally observed apparent diffusion constant 0.01 *µm*^2^*/s*, we get *τ*_*sim*_ ∼1 second.

A different approach to calibration is to use an apparent viscosity of the nucleoplasm. Using the viscosity of water *η* = 0.001 Pa-s, which is definitely an underestimate, we can write the drag coefficient: *γ* = 6*πησ*. Equating this with the previously mentioned definition of *γ*, we get the underestimated value *τ*_*sim*_ ∼0.01 second.

Combining both these approaches, we use an order of magnitude calibration: *τ*_*sim*_ = 0.1 seconds.

### 4. Polymer potential

The polymer potential constitutes of two terms: bonding between nearest neighbors and a short-range inter-monomer repulsion to simulate self-avoidance. The nearest neighbors along the polymer chain are bonded using a linear spring: *U*_*nn*_ = (*k/*2)(*r*−*d*)^2^, where *k* is the spring constant and *d* is the unperturbed bond length. We use *k* = 30*ϵ/σ*^2^ unless otherwise mentioned. Here *ϵ* is the reduced unit of energy. For Rouse chains we used *d* = 0 while for all else we set *d* = 1*σ*.

Self avoidance is modeled using a pairwise soft-core repulsive potential of the form: *U*_*sa*_(*r*) = (*E*_*0*_*/*2)(1 + tanh(1 − *k*_*sa*_(*r*−*d*_*sa*_))), where *k*_*sa*_ = 20*ϵ/σ*^2^ encodes the steepness of the repulsion, *d*_*sa*_ = 1*σ* is typical distance below which repulsion kicks in, and *E*_*0*_ = 5*ϵ* is the maximum repulsive energy.

Finally, there is the spherical-confinement potential. We use a flat-bottom harmonic or half-harmonic restraint for the confinement: *U*_*conf*_ = (*k*_*conf*_ */*2)(*r* − *R*_*c*_)^2^Θ(*r*−*R*_*c*_), where Θ(*r*−*R*_*c*_) is the Heavy-side theta function that is zero when *r < R*_*c*_ and one when *r* ≥ *R*_*c*_. We use *k*_*conf*_ = 30*ϵ/σ*^2^ throughout this work.

#### a. Chromosome potential

For the chromosome potential We utilize the optimized MiChroM parameters [19, 61] to simulate the passive forces within our Brownian dynamics scheme (Eq. 1). We chose chromosome 10 of Human GM12878 cell line, which has *N* = 2711 beads with each bead representing 50 kb DNA. The active loci were selected to be at the monomer indices: 99,376,740,1100,1432,1860,2340. We then studied the effect on the passive structure as the activity was turned on for the active loci.

### 5. Simulation trajectory analysis

Starting from random configurations, the polymer is equilibrated at temperature *T*. We used *k*_*B*_*T* = 0.2*ϵ* for simulations unless otherwise mentioned. Then, activity with a fixed *F* and *τ* is turned on for every monomer, and simulations are run for 10^6^ time steps with integration step *dt* = 10^*−*3^*τ*_*sim*_ to establish the active steady state. We also used the analytical calculations of the

Rouse polymer as a verification step for our simulations (Supplementary Materials). Steady-state simulation for each parameter set was run for 10^4^*τ*_*sim*_, saving the particle coordinates every 0.1*τ*_*sim*_. We simulated multiple replicas for each parameter for statistical analyses. All reported quantities, like the radius of gyration, monomer density, and MSD, were computed from the steady-state trajectories. These quantities are also averaged over the ensemble.

#### a. Monomer density

The monomer density *n*(*r*) is computed by counting the number of monomers in concentric shells, such that: 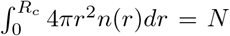, where *N* is the total number of monomers.

#### b. Radial distribution function

The radial distribution function is computed from the standard definition: *g(r)* = *dn*_*r*_*σ*^3^*/*(4*πr*^2^*drϕ*), where *dn*_*r*_ is the number of monomers within distance *r* and *r* + *dr* from the center.

#### c. Displacement correlation between particles

The spatial correlation of displacements between particles was computed using the formula: 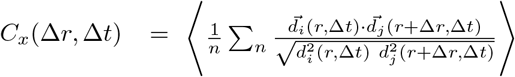, where *d*_*i*_(*r*, Δ*t*) is the displacement of the *i*-th particle located at *r* over a lag time Δ*t* and *d*_*j*_ is the corresponding displacement of the *j*-th particle that is located a distance Δ*r* away from the *i*-the particle. The displacement correlation is summed over for all the *n* particles that lie within a sphere of radius Δ*r*, and the angular bracket denotes averaging over the trajectories.

#### d. Phase-separation coefficient

The phase-separation coefficient *µ* is defined as: *µ* ≡(⟨*n*_*AA*_⟩ + ⟨*n*_*PP*_⟩)*/* ⟨2*n*_*AP*_⟩. Here, *n*_*AA*_ is the number of pairwise contacts between *A* blocks (homotypic contacts), and *n*_*P P*_ (homotypic) and *n*_*AP*_ (heterotypic) are similarly defined. Two monomers are defined to be in contact when the distance between them is less than 1.5*σ*. Contacts between monomers of the same block are ignored to enhance the inter-block phase separation signal.

#### e. Active regime diagrams

Steady-state trajectories were generated for a logarithmically spaced grid with *F* varying between 0 − 15*ϵ/σ* and *τ* varying between 0.1 − 300*τ*_*sim*_. The observables, like the radius of gyration, were computed for the above-mentioned grid and then interpolated into a finer grid to plot the active regime diagrams (Figs. 2A, 3B).

#### f. Contact maps

The contact maps were generated using the procedure described in Refs. [19, 61]. The pairwise distance between monomers *r*_*ij*_ is converted to probability using the previously optimized sigmoid function *f*_*contact*_(*r*_*ij*_) = 0.5 ∗ (1 + tanh(3.22 ∗ (1.78− *r*_*ij*_))) [19, 61].

A tutorial on how to run the active simulations can be found at https://github.com/s-brahmachari/ActiveOpenMiChroM/tree/main/Tutorials/Active_Chromosome_Dynamics. The simulation snapshots were generated using the Visual Molecular Dynamics (VMD) software http://www.ks.uiuc.edu/Research/vmd/.

## Appendix B: Active Rouse polymer

Rouse chains, both because of their simplicity and analytical tractability, are a paradigm for studying polymer structure and dynamics. Considering a chain of *N* monomers, the equation of motion for the *n*-hth Rouse monomer reads as follows [41, 43].

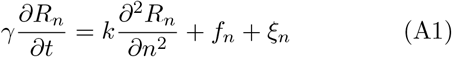

where *k* is the spring constant associated with the harmonic bonds connecting the neighboring monomers.

### 1. Motor activity characterizes the relaxation spectrum of Rouse modes

Defining the *p*-th normal mode of the Rouse chain, representing the dynamics in a subchain of size *N/p*, as follows.

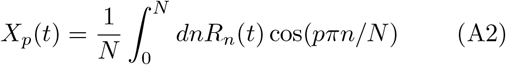

We obtain the autocorrelation between the Rouse modes for active chains [41]:

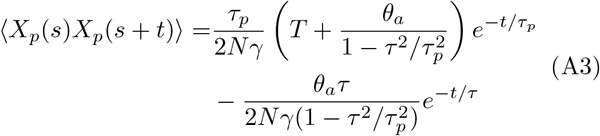

where *τ*_*p*_ = *N* ^2^*γ/*(*kπ*^2^*p*^2^) is the equilibrium relaxation time of the *p*-th mode. The longest Rouse relaxation time is given by *τ*_Rse_ = *N* ^2^*γ/*(*kπ*^2^).

Figure A1A-B shows the autocorrelation of Rouse modes of the active and passive chains from simulations and theory (Eq. (A3)). The Rouse mode *p* with an equilibrium relaxation time *τ*_*p*_ that is longer than the active correlation time (*τ*_*p*_ ≫ *τ*), appears equilibrium-like, i.e., the autocorrelation decays exponentially with a time-constant ≈*τ*_*p*_ (Fig. A1B). However, activity raises the effective temperature associated with the modes to *T* + *θ*_*a*_. On the other hand, when the equilibrium relaxation time of the Rouse mode *p* is shorter than the activity persistent time (*τ*_*p*_ ≪*τ*), the mode autocorrelation relaxes exponentially with a time constant dominated by the active correlation time *τ*. This leads to crowding of the relaxation curves near the active correlation time in Fig. A1B.

### 2. Motor-induced persistent activity leads to diverse dynamic regimes

The scaling of mean-squared displacement (MSD) with time (*t*): ∼*t*^α^, defines the dynamic regime of the beads, such as diffusion (*α* = 1), sub-diffusion (*α <* 1), and super-diffusion (*α >* 1) which includes ballistic motion (*α* = 2). The MSD for the center-of-mass (CM) of a chain, governed by the zero wavenumber mode: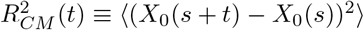, is given by,

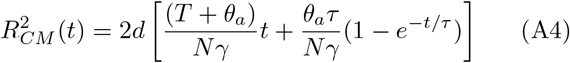

where *d* = 3 is the dimension of space coordinates. The first term within brackets, proportional to *t*, is the diffusive term that dominates at long times (*t* ≫ *τ*). Note, at long times, the diffusion constant-like coefficient associated with the active polymer CM motion is 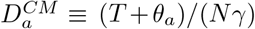. Hence, at long times, the CM motion appears diffusive with an effective temperature *T* + *θ*_*a*_ and the expected drag coefficient of a Rouse polymer *Nγ*. While at shorter times, there is a leading order super-diffusive (ballistic) term (∼ *t*^2^) that dominates thermal diffusion: 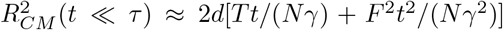 (Fig. A1C).

**FIG. A1.**
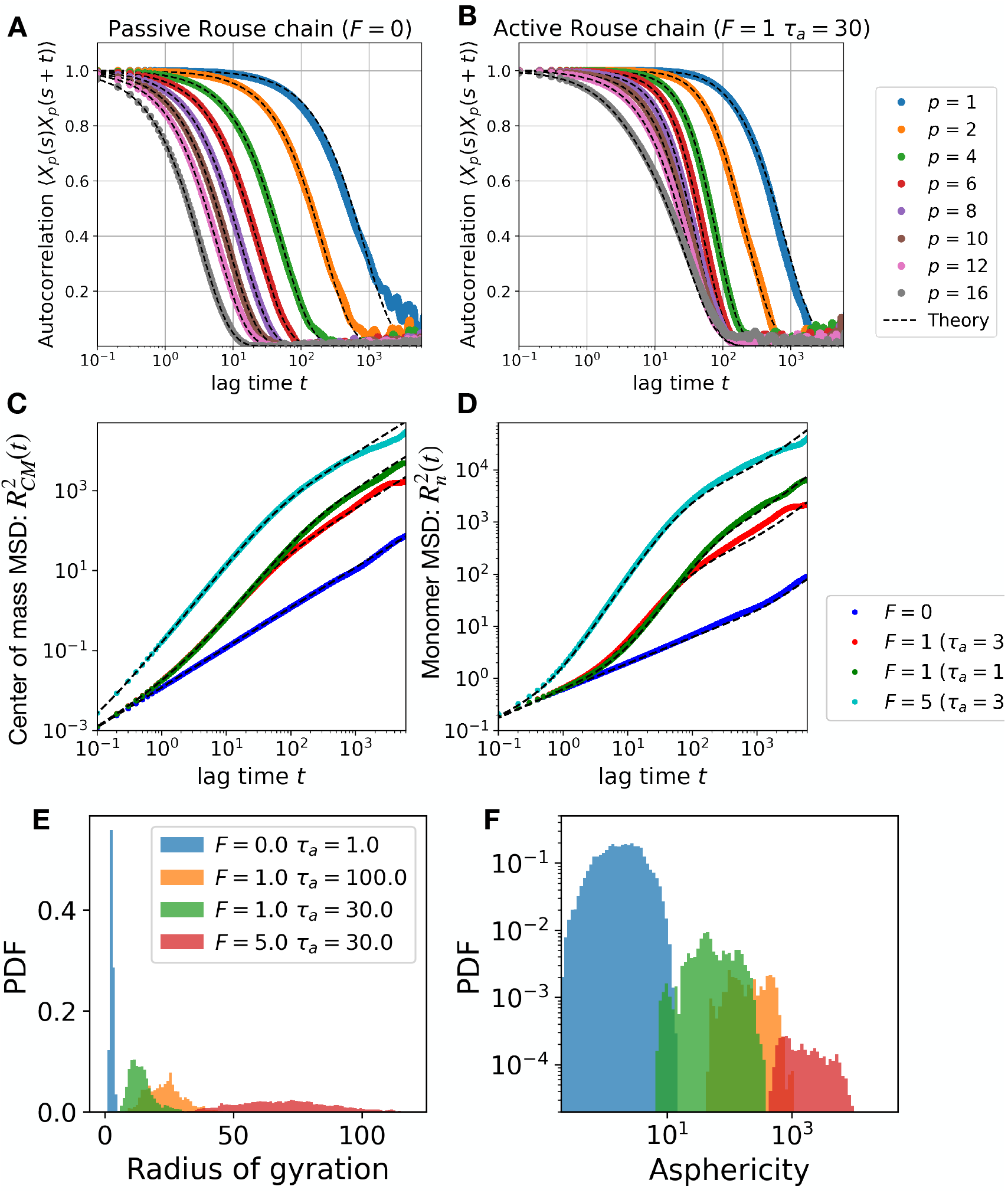
Free Rouse chain. Rouse chain of size *N* = 500 and spring constant *k* = 30*ϵ/σ*^2^ simulated at *k*_**B**_ *T* = 1. Autocorrelation of Rouse modes for (A) Passive and (B) active chains. The different colors represent different wavemodes as shown in the legend. Note that for active chains the modes that relax at times longer than the correlation time *τ* = 30 show equilibrium-like behavior. While the modes with characteristic relaxation time smaller than the correlation time have a prolonged relaxation under the active noise. The dashed lines show analytical results from Eq. (A3). Mean-squared displacements for (C) center-of-mass motion and (D) monomer motion for Rouse chains. The longest Rouse relaxation time for this polymer is *τ*_*1*_ ≈ 850. The dashed lines show theoretical expectations from Eqs. (A4) and (A5). (E) Radius of gyration of Rouse polymer for various values of *F* and *τ*. Increasing activity *θ*_**a**_ leads to a higher radius of gyration. (F) Distributions of asphericity of the polymer configurations, derived from the eigenvalues of the gyration tensor, show anisotropic and spheroidal structures with increasing activity.

On the other hand, the MSD for the *n*-th monomer is given by a sum over all the Rouse modes *p*:

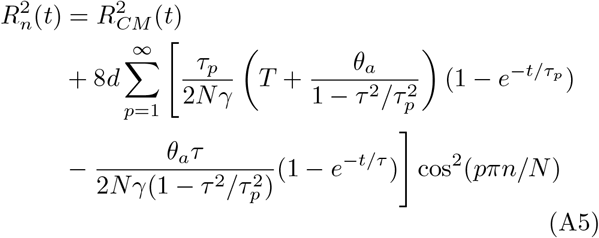

where 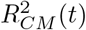 (*t*) is defined in Eq. (A4), and *d* = 3 is the space dimension. Firstly, note that at very long times, when all the Rouse modes have relaxed (*t* ≫ *τ*_*p*_ and *t*≫ *τ*), monomer dynamics is governed by the CM motion, given by Eq. (A4). This transition occurs at lag times longer than the longest Rouse time *τ*_*1*_ (Fig. A1D).

Interesting dynamic behaviors are observed at times comparable to the activity persistent time (*τ*) and *τ*_Rse_. In absence of activity, i.e., a passive system (*θ*_*a*_ = 0), monomer MSD shows subdiffusion in the regime of lag times smaller than *τ*_Rse_, while it shows diffusion corresponding to CM motion for lag times larger than *τ*_Rse_ (Fig. A1). For an active system, there is super-diffusive (near ballistic) signatures in the monomer MSD at times smaller or comparable to the active persistent time *τ*. This is a direct consequence of correlated noise, that consistently propels the particles in a specific direction before changing the direction at lag times comparable to the persistent time. For lag times beyond the active persistent time but smaller than the Rouse time (*τ*_Rse_), the monomers move subdiffusively, albeit with a higher effective temperature (Fig. A1).

### 3. Motor activity drives swelling of Rouse polymer

Motor activity not only influences the dynamics, it is consequential for the conformations as well. Motor-generated persistent forces stretch the polymer, leading to swollen conformations that exhibit a larger radius of gyration (Fig. A1E). The temperature-like quantity associated with activity: *θ*_*a*_ ≡ *F* ^2^*τ/γ*, when dominates the thermal temperature *T < θ*_*a*_, there is activity-driven swelling of the polymer. The swollen active polymers are spheroidal, where asphericity increases with the active temperature *θ*_*a*_ (Fig. A1F).

### 4. Confined Rouse polymer

Rouse chain in confinement does not show the structural regime corresponding to collapse at the center (Fig. A2). This is because self-avoidance is a necessary ingredient for center collapse. The monotonically increasing radius of gyration for Rouse polymers leads to only boundary enrichment when the mean persistent path is longer than the confinement dimension.

## Appendix C: Chromosome model

We use the optimized MiChroM chromosome potential as described in Refs. [19, 61] to simulate chromosome 10 of the Human GM12878 lymphoblastoid cell using a 50 kb resolution model (*N* = 2712). MiChroM uses structural annotations to assign A and B-type beads to the chromosome polymer and then the interaction profiles between the monomer types lead to the plaid-like patterns of interactions as seen in the Hi-C maps. There is an additional ideal chromosome term that drives length-wise compaction or crumpling of the polymer along the contour, which is a feature of SMC-driven compaction. The simulated MiChroM contact maps and the corresponding experimental hic maps [13] are shown in Fig. A10.

We added seven active loci along the chromosome at monomer indices (99,376,740,1100,1432,1860,2340) and varied the activity in those loci to observe the corresponding change in the structure. The contact maps and representative structures for the passive case (*F* = 0) and active steady state with (*F* = 3, *τ* = 30) and (*F* = 5, *τ* = 1), simulated at *k*_*B*_*T* = 1, are shown in Fig. A11. Long correlation times representing persistent transcription activity drive the active loci to the periphery of chromosome territories.

**FIG. A2.**
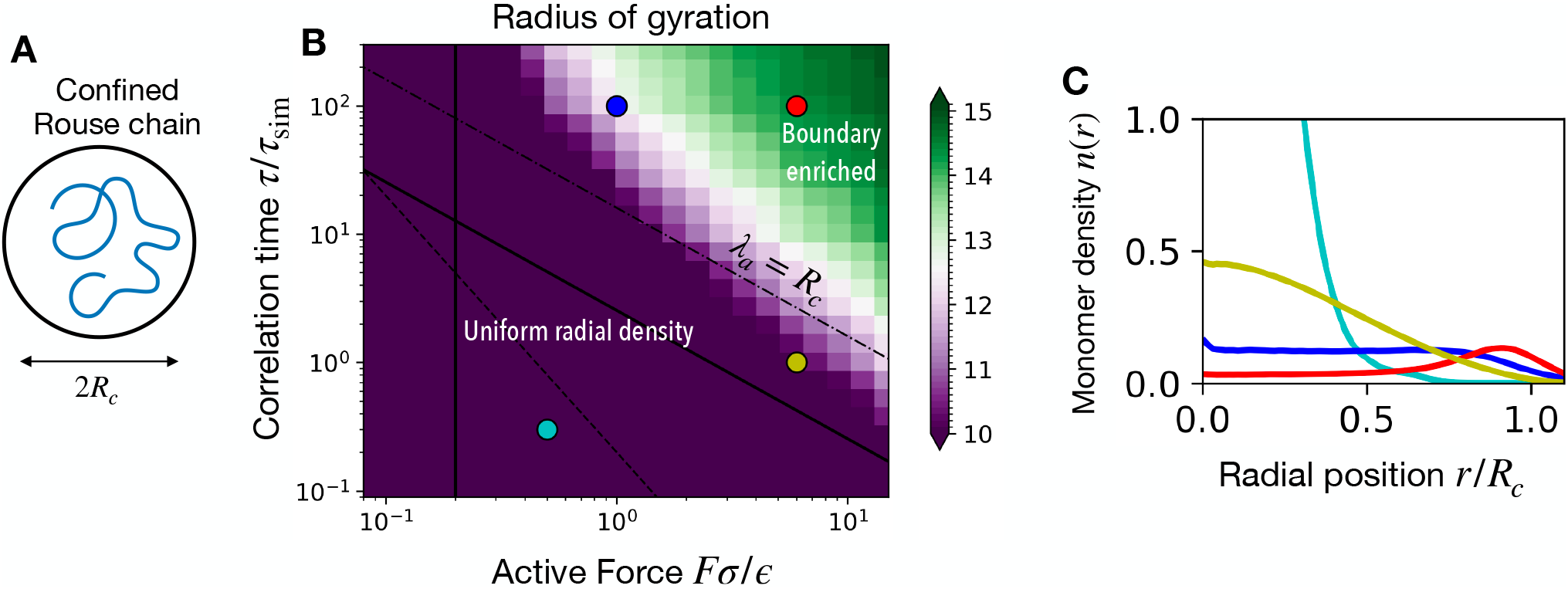
Confined Rouse chain. Rouse polymer of size *N* = 2000 in spherical confinement of size *R*_*0*_ = 16*σ*. (A) Active regime diagram showing the radius of gyration as a function of the active force and correlation time. Boundary enrichment occurs when the mean persistent path *λ*_**a**_ exceeds the confinement dimension *R*_**c**_. (C) Radial monomer density profile shows a monotonic decrease of density at the center with increasing activity, while the boundary becomes increasingly enriched.

**FIG. A3.**
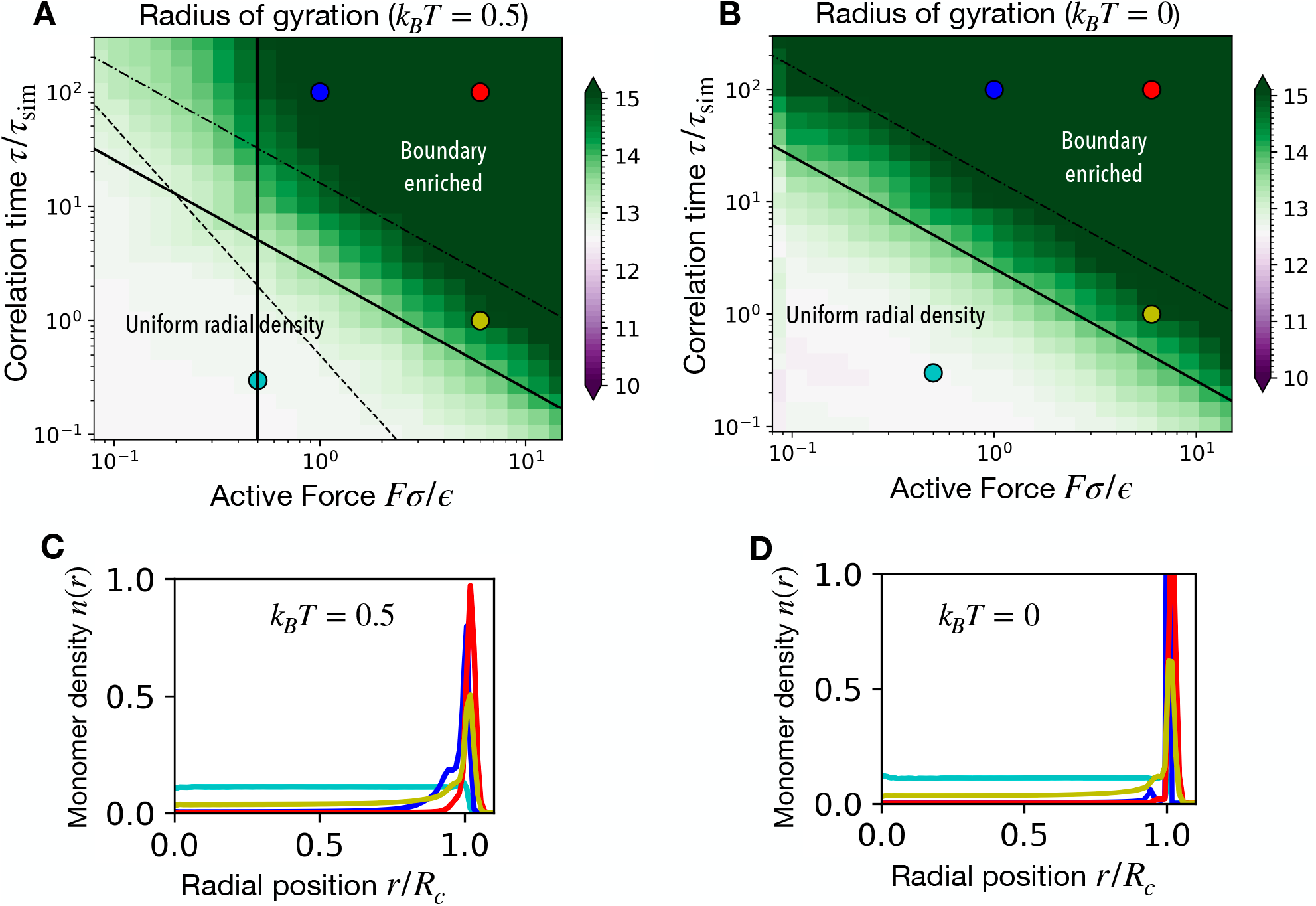
Self-avoiding gas. *N* = 2000 self-avoiding active particles confined within a spherical cavity of radius *R*_*0*_ = 16*σ* simulated at two temperatures: *k*_**B**_ *T* = 0 and *k*_**B**_ *T* = 0.5. (A) and (B) Active regime diagram showing the radius of gyration of the gas particles s a function of the active force and the correlation time. There is a monotonic increase in the radius of gyration for high activity. (C) and (D) Monomer density *n*(*r*) variation with the radial position. The boundary-enriched state may be observed when the mean persistent path is higher than the confinement dimension.

**FIG. A4.**
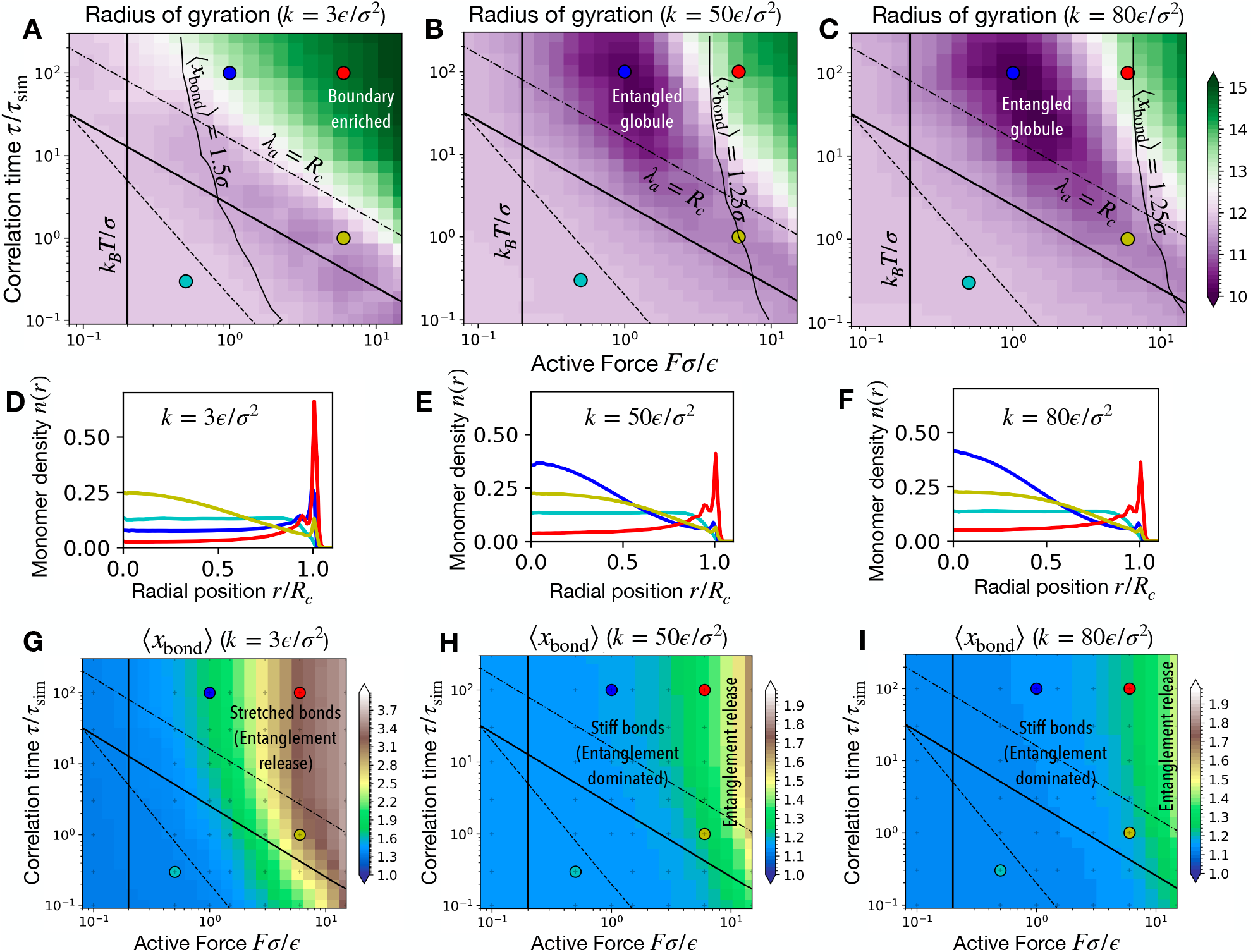
Effect of spring constant on the entangled globule state. Self-avoiding polymer of size *N* = 2000 in spherical confinement with radius *R*_*0*_ = 16*σ* with various values of the spring constant *k*. (A)-(C) Active regime diagrams plotting the radius of gyration show the enhanced stability of the entangled globule state for higher spring constants *k*. Note the shift of the mean bond length line (⟨*x*_bond_⟩ = 1.25*σ*) along with the boundary of the entangled globule. (D)-(F) Radial profile of monomer density for various spring constants. (G)-(I) Contour plot showing the average bond length as a function of active force and correlation time for various spring constants. When the bond length is higher than 1.25*σ* the topological constraints are diminished leading to entanglement release and destabilization of the entangled globule state.

**FIG. A5.**
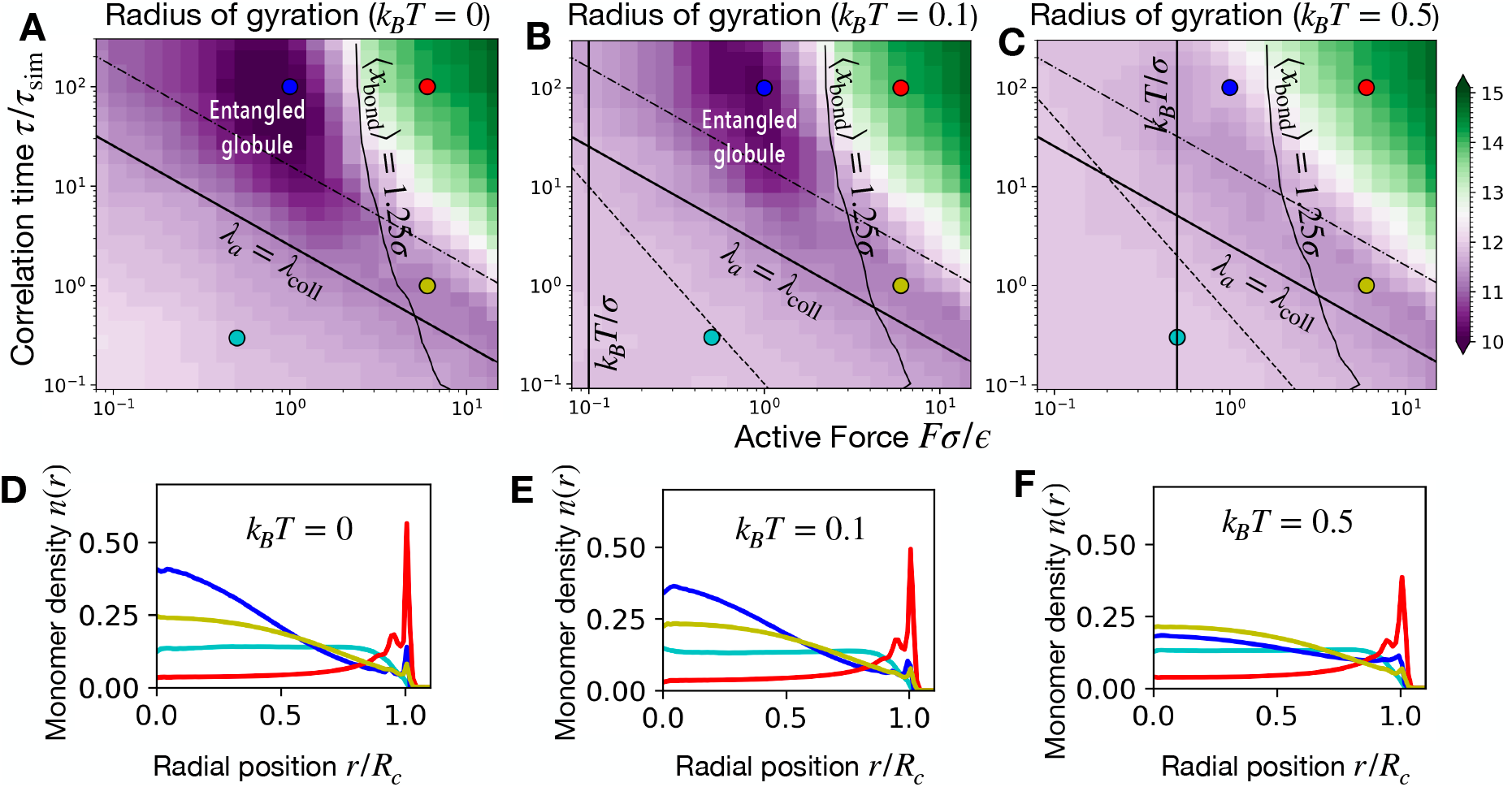
Effect of temperature in destabilizing the entangled globule state. Self-avoiding polymer of size *N* = 2000 in spherical confinement with radius *R*_*0*_ = 16*σ* simulated at various temperatures *T*. (A)-(C) Active regime diagrams plotting the radius of gyration show the enhanced stability of the entangled globule state for lower temperatures. Note the shift of the thermal amplitude line *k*_**B**_ *T/σ*. (D)-(F) Radial profile of monomer density for various spring constants. When the thermal noise amplitude dominates the active noise, *F < k*_**B**_ *T/σ* the active correlations are destroyed and the entangled globule state is destabilized.

**FIG. A6.**
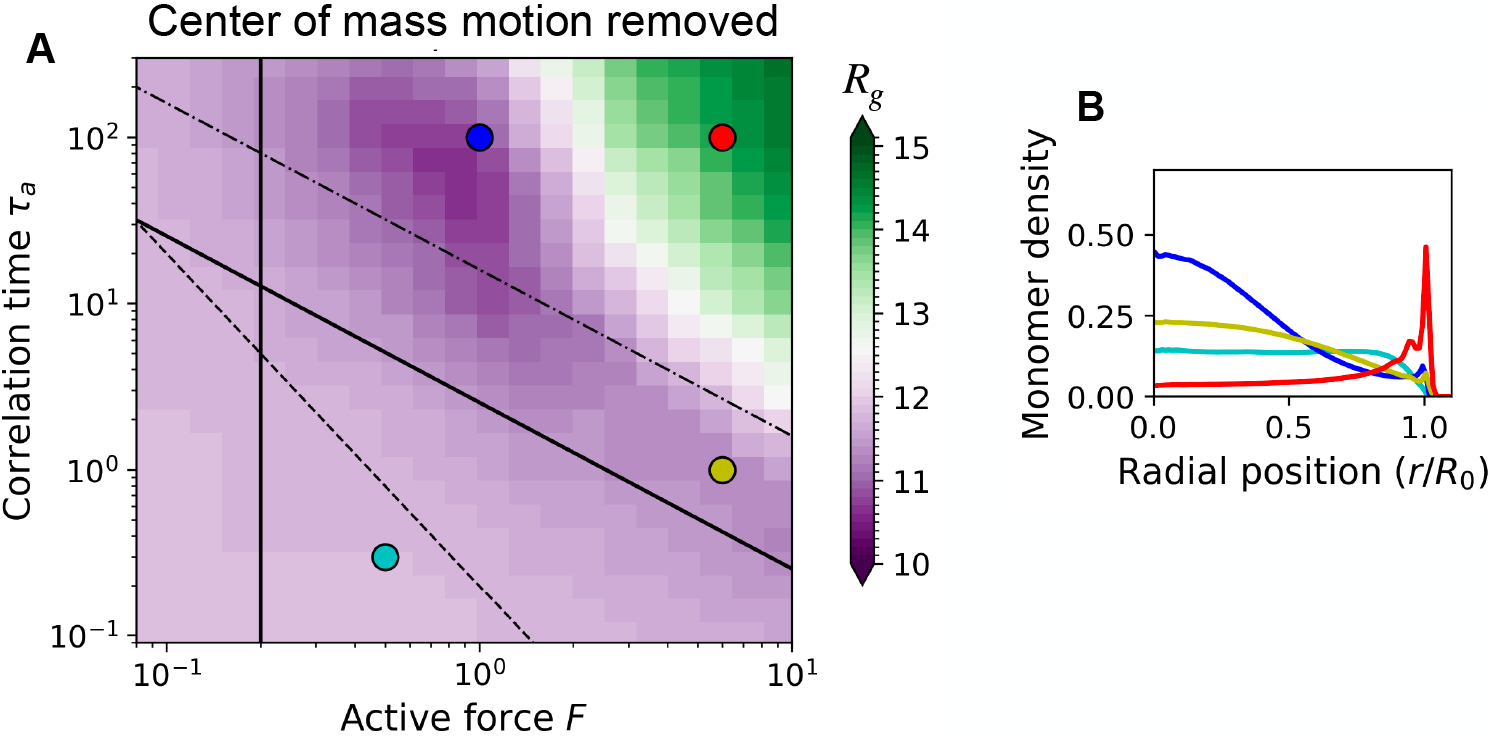
Effect of the explicit center of mass motion correction on the active polymer regimes. (A) The radius of gyration of the active polymer for various active forces and correlation times is calculated in simulations where the active noise for each monomer is corrected so that the net center of mass motion is reduced. We implemented this using the CMMotionRemover scheme in OpenMM [101]. (B) Monomer radial density for the various regimes shown in the panel (A). As can be noted by comparison with Figs. 2A-B, the center of mass motion correction does not affect the active polymer regimes. This is a consequence of the active forces being balanced due to their random orientation (see Discussion).

**FIG. A7.**
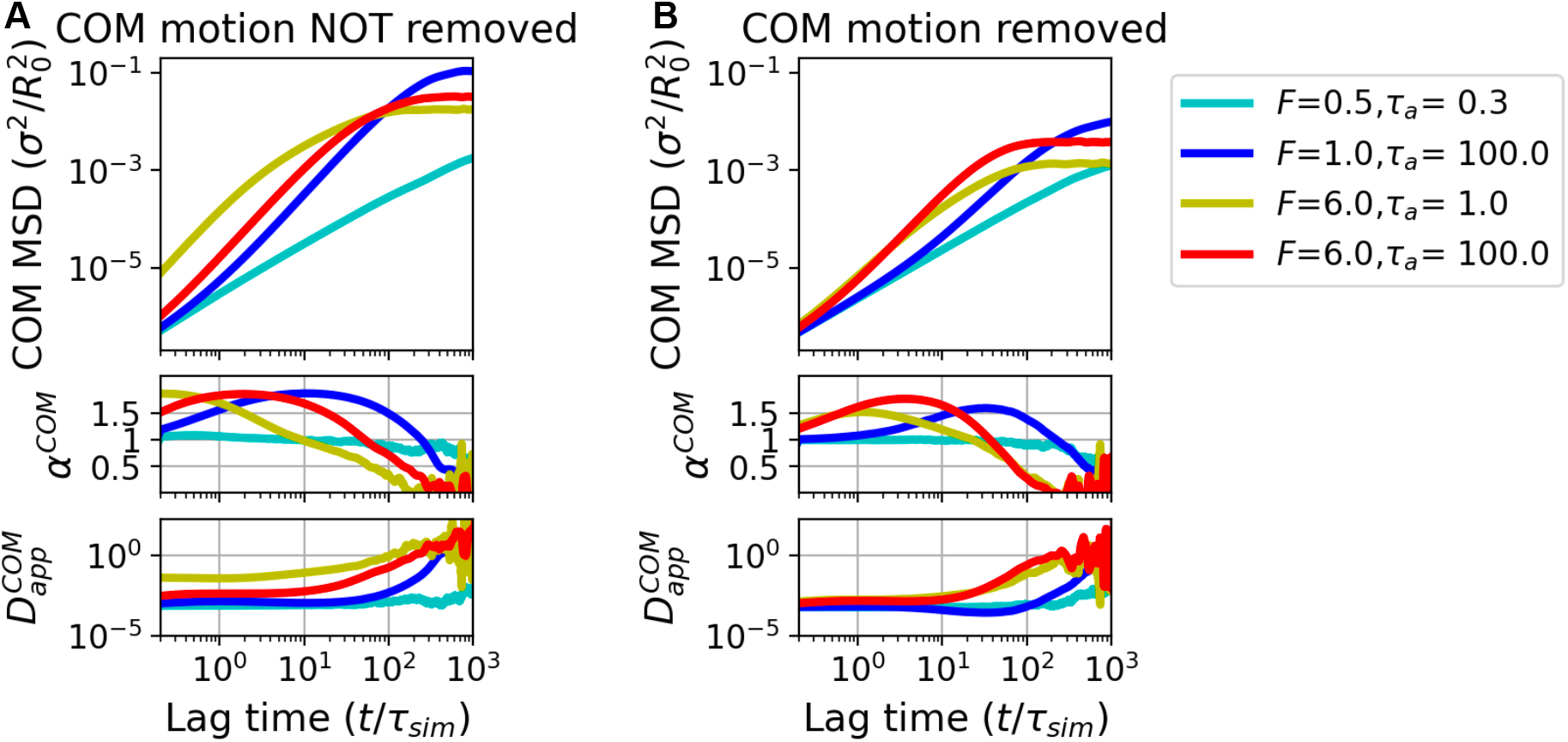
Effect of the explicit center of mass motion correction on the center of mass mean-squared displacement. (A) MSD of the center of mass (COM) of the active polymer without any motion correction for the center of mass. The subsequent panels show the apparent MSD exponent and the apparent diffusion constant. These simulations correspond to the ones shown in Fig. 2. (B) Same as in (A) but with COM motion correction implemented. Note, as expected, that the MSD and the apparent COM diffusion constant are suppressed upon COM motion removal.

**FIG. A8.**
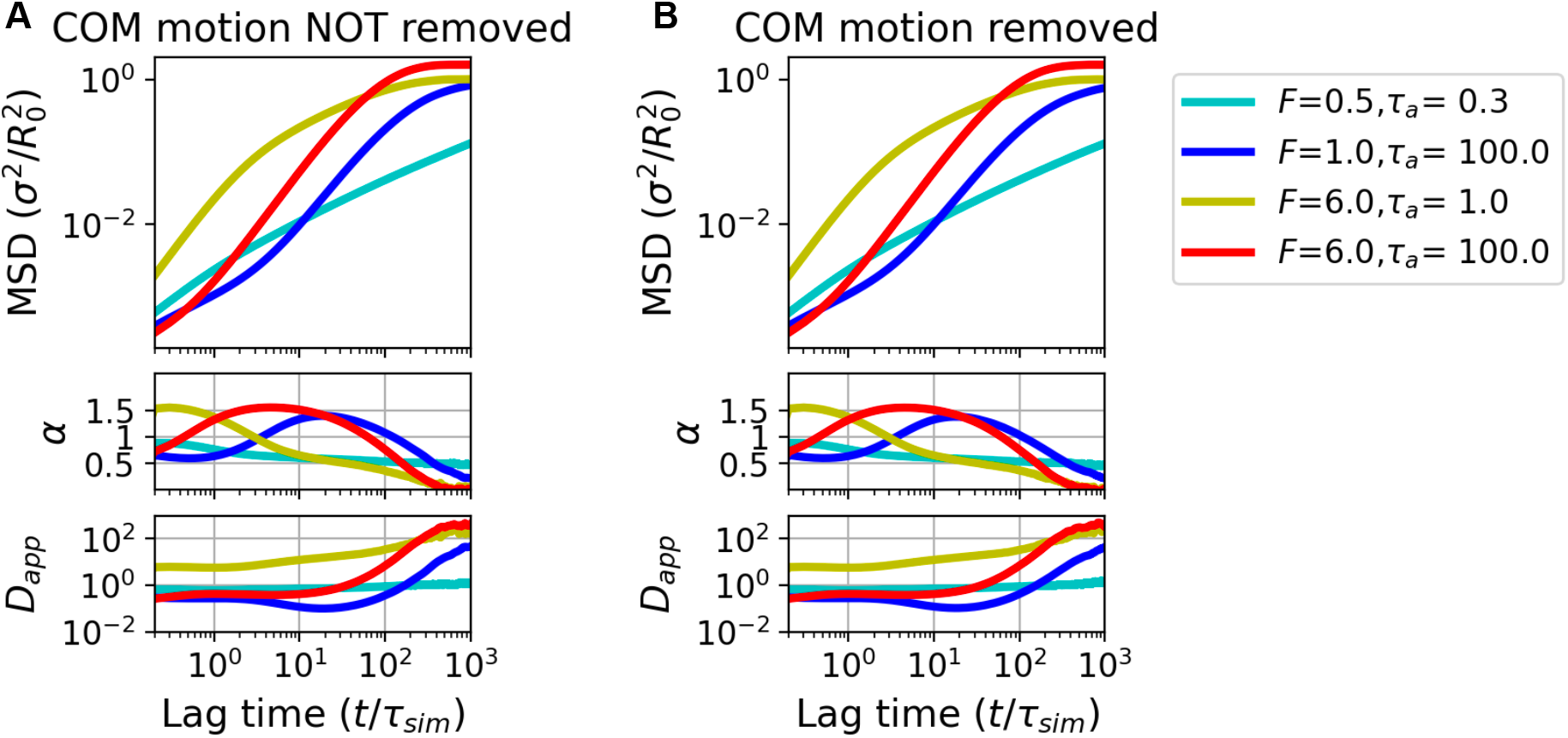
Effect of the explicit center of mass motion correction on the monomer mean-squared displacement. Average MSD of a monomer, the MSD exponent, and the apparent diffusion constant, without (A) and with (B) COM motion correction. Panel (A) is the same as in Fig. 2F. Note that the monomer MSDs are unchanged upon COM motion removal, which again is a direct consequence of the active forces already being balanced due to their random orientation (see Discussion).

**FIG. A9.**
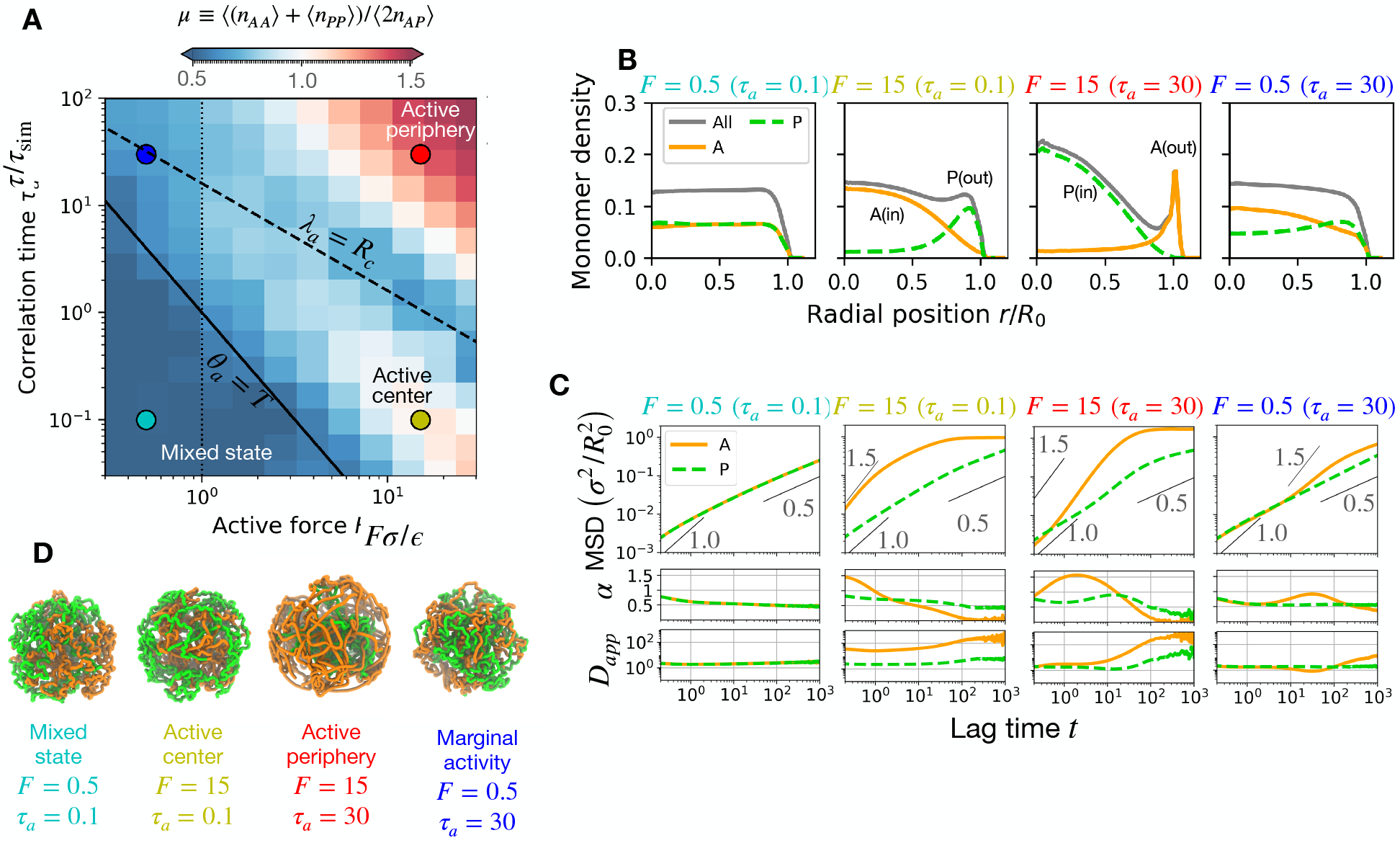
Confined active-passive block copolymer at higher thermal temperature. (A) Active regime diagram of block copolymer in confinement at thermal temperature *k*_**B**_ *T* = 1.0*ϵ*. Higher temperatures destabilize the entangled globule leading to better segregation of active and passive blocks. Representative regions are selected on the *F* − *τ* phase space to show in the corresponding subfigures. (B) Radial monomer density of active and passive blocks showing radial segregation. (C) MSD versus lag time. MSD exponent and apparent diffusivity are shown in subpanels. (D) Simulation snapshots of the block copolymer.

**FIG. A10.**
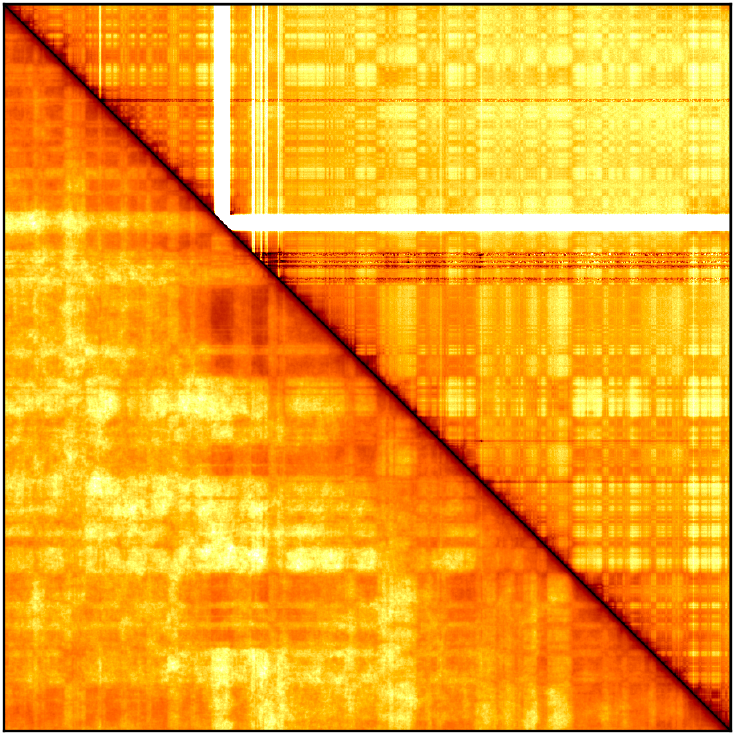
Comparison with experiments. Comparing Hi-C maps from Rao *et al*. [13] with the base MiChroM model [19, 61], obtained by setting activity to zero (*F* = 0) and *k*_**B**_ *T* = 1.0*ϵ*. The upper triangle shows experimental data and the lower triangle is the simulation result for chromosome 10 of the Human GM12878 cell line at 50 kb resolution.

**FIG. A11.**
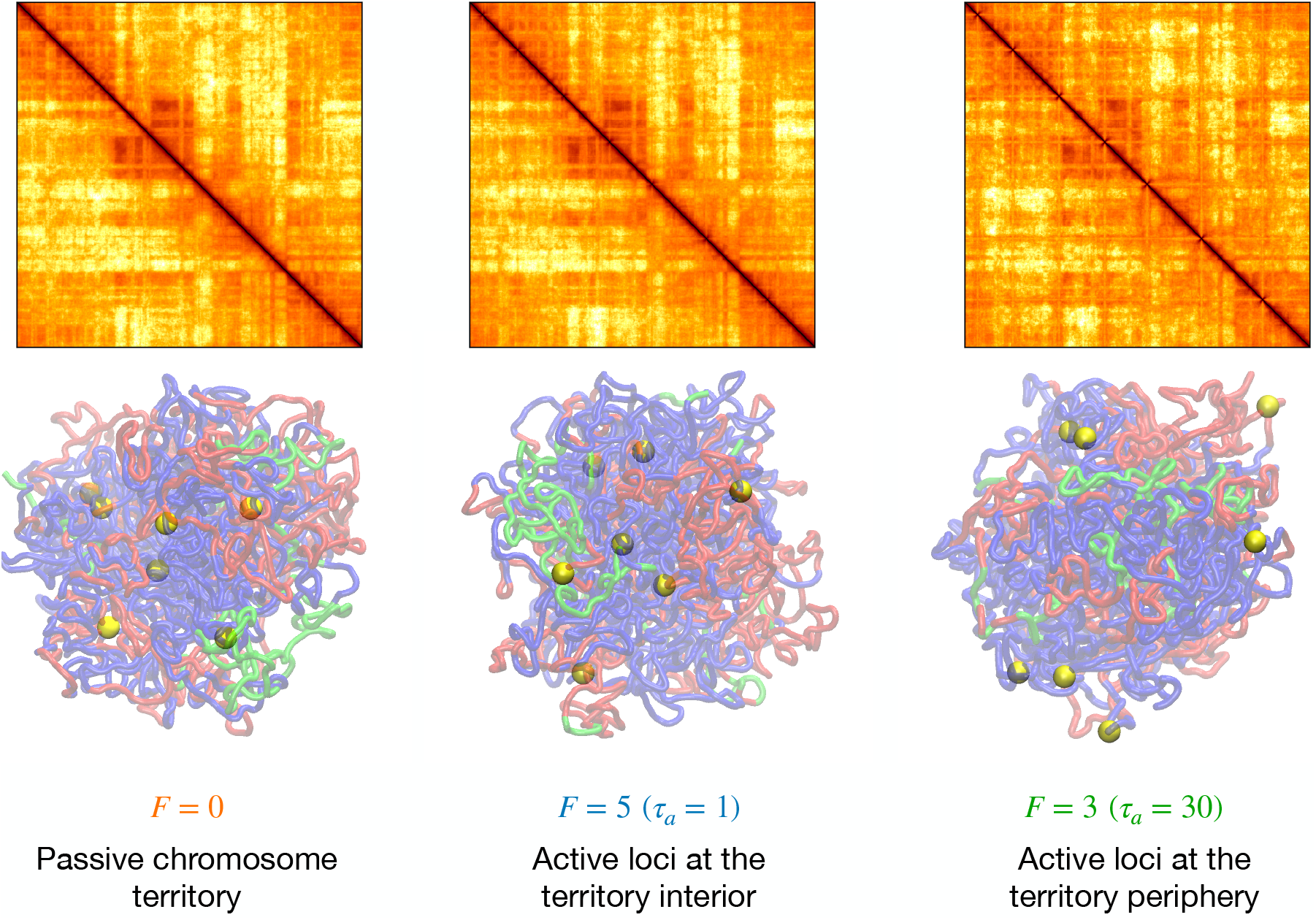
Active perturbations to GM12878 chromosome 10. Simulation contact maps and snapshots of the GM12878 chromosome 10 for the passive and active cases. In the snapshots, *A* type and *B* type beads are colored in red and blue respectively. In the active scenario, 7 discrete loci, shown as yellow spheres, have active noise while all the rest just have MiChroM potential. Active loci are more dynamic that perturb the compartments locally. Active loci with a long correlation time move to the periphery of the chromosome territory. The overall effect on the contact maps is minimal due to the low density of active loci, which only perturbs the effective-equilibrium structure locally.

## References

[1] D. Mizuno, C. Tardin, C. F. Schmidt, and F. C. MacK-intosh, Nonequilibrium mechanics of active cytoskeletal networks, Science 315, 370 (2007).

[2] P. M. Bendix, G. H. Koenderink, D. Cuvelier, Z. Dogic, B. N. Koeleman, W. M. Brieher, C. M. Field, L. Mahadevan, and D. A. Weitz, A quantitative analysis of contractility in active cytoskeletal protein networks, Biophysical Journal 94, 3126 (2008).

[3] S. Wang and P. G. Wolynes, Active contractility in actomyosin networks, Proceedings of the National Academy of Sciences of the United States of America 109, 6446 (2012).

[4] S. Wang and P. G. Wolynes, On the spontaneous collective motion of active matter, Proceedings of the National Academy of Sciences 108, 15184 (2011), 10.1073/pnas.1112034108.

[5] E. Tjhung, D. Marenduzzo, and M. E. Cates, Spontaneous symmetry breaking in active droplets provides a generic route to motility, Proceedings of the National Academy of Sciences of the United States of America 109, 12381 (2012).

[6] L. Blanchoin, R. Boujemaa-Paterski, C. Sykes, and J. Plastino, Actin dynamics, architecture, and mechanics in cell motility, Physiological Reviews 94, 235 (2014).

[7] V. Hakim and P. Silberzan, Collective cell migration: a physics perspective, Reports on Progress in Physics 80, 076601 (2017).

[8] T. Cremer and C. Cremer, Chromosome territories, nuclear architecture and gene regulation in mammalian cells, Nature Reviews Genetics 2001 2:4 2, 292 (2001).

[9] W. A. Bickmore, The spatial organization of the human genome 10.1146/annurev-genom-091212-153515 (2013).

[10] C. Hoencamp, O. Dudchenko, A. M. Elbatsh, S. Brahmachari, J. A. Raaijmakers, T. van Schaik, Ángela Sedeño Cacciatore, V. G. Contessoto, R. G. van Heesbeen, B. van den Broek, A. N. Mhaskar, H. Teunissen, B. G. S. Hilaire, D. Weisz, A. D. Omer, M. Pham, Z. Colaric, Z. Yang, S. S. Rao, N. Mitra, C. Lui, W. Yao, R. Khan, L. L. Moroz, A. Kohn, J. S. Leger, A. Mena, K. Holcroft, M. C. Gambetta, F. Lim, E. Farley, N. Stein, A. Haddad, D. Chauss, A. S. Mutlu, M. C. Wang, N. D. Young, E. Hildebrandt, H. H. Cheng, C. J. Knight, T. L. Burnham, K. A. Hovel, A. J. Beel, P. J. Mattei, R. D. Kornberg, W. C. Warren, G. Cary, J.L. Gómez-Skarmeta, V. Hinman, K. Lindblad-Toh, F. D. Palma, K. Maeshima, A. S. Multani, S. Pathak, L. Nel-Themaat, R. R. Behringer, P. Kaur, R. H. Medema, B. van Steensel, E. de Wit, J. N. Onuchic, M. D. Pierro, E. L. Aiden, and B. D. Rowland, 3d genomics across the tree of life reveals condensin ii as a determinant of architecture type, Science (New York, N.Y.) 372, 10.1126/SCIENCE.ABE2218 (2021).

[11] E. Lieberman-aiden, N. L. V. Berkum, L. Williams, M. Imakaev, T. Ragoczy, A. Telling, I. Amit, B. R. Lajoie, P. J. Sabo, M. O. Dorschner, R. Sandstrom, B. Bernstein, M. A. Bender, M. Groudine, A. Gnirke, J. Stamatoyannopoulos, and L. A. Mirny, Comprehensive mapping of long-range interactions reveals folding principles of the human genome, Science 33292, 289 (2009).

[12] J. R. Dixon, I. Jung, S. Selvaraj, Y. Shen, J. E. Antosiewicz-Bourget, A. Y. Lee, Z. Ye, A. Kim, N. Rajagopal, W. Xie, Y. Diao, J. Liang, H. Zhao, V. V. Lobanenkov, J. R. Ecker, J. A. Thomson, and B. Ren, Chromatin architecture reorganization during stem cell differentiation, Nature 518, 331 (2015).

[13] S. S. P. Rao, M. H. Huntley, N. C. Durand, E. K. Stamenova, I. D. Bochkov, J. T. Robinson, A. L. Sanborn, I. Machol, A. D. Omer, E. S. Lander, and E. Lieberman-Aiden, Article a 3d map of the human genome at kilobase resolution reveals principles of chromatin looping, Cell 159, 1665 (2014).

[14] A. N. Boettiger, B. Bintu, J. R. Moffitt, S. Wang, B. J. Beliveau, G. Fudenberg, M. Imakaev, L. A. Mirny, C. T. Wu, and X. Zhuang, Super-resolution imaging reveals distinct chromatin folding for different epigenetic states, Nature 2016 529:7586 529, 418 (2016).

[15] B. Bintu, L. J. Mateo, J.-H. Su, N. A. Sinnott-Armstrong, M. Parker, S. Kinrot, K. Yamaya, A. N. Boettiger, and X. Zhuang, Super-resolution chromatin tracing reveals domains and cooperative interactions in single cells, Science 362, eaau1783 (2018).

[16] E. M. Hildebrand and J. Dekker, Mechanisms and functions of chromosome compartmentalization, Trends in biochemical sciences 45, 385 (2020).

[17] J. H. Su, P. Zheng, S. S. Kinrot, B. Bintu, and X. Zhuang, Genome-scale imaging of the 3d organization and transcriptional activity of chromatin, Cell 182, 1641 (2020).

[18] M. Falk, Y. Feodorova, N. Naumova, M. Imakaev, B. R. Lajoie, H. Leonhardt, B. Joffe, J. Dekker, G. Fudenberg, I. Solovei, and L. A. Mirny, Heterochromatin drives compartmentalization of inverted and conventional nuclei, Nature 2019 570:7761 570, 395 (2019).

[19] M. D. Pierro, B. Zhang, E. L. Aiden, P. G. Wolynes, and J. N. Onuchic, Transferable model for chromosome architecture, Proceedings of the National Academy of Sciences of the United States of America 113, 12168 (2016).

[20] Y. Qi, A. Reyes, S. E. Johnstone, M. J. Aryee, B. E. Bernstein, and B. Zhang, Data-driven polymer model for mechanistic exploration of diploid genome organization, Biophysical Journal 119, 1905 (2020).

[21] A. Esposito, S. Bianco, A. M. Chiariello, A. Abraham, L. Fiorillo, M. Conte, R. Campanile, and M. Nicodemi, Polymer physics reveals a combinatorial code linking 3d chromatin architecture to 1d chromatin states, Cell Reports 38, 10.1016/J.CELREP.2022.110601 (2022).

[22] H. Wong, H. Marie-Nelly, S. Herbert, P. Carrivain, H. Blanc, R. Koszul, E. Fabre, and C. Zimmer, A predictive computational model of the dynamic 3d interphase yeast nucleus, Current Biology 22, 1881 (2012).

[23] D. Jost, P. Carrivain, G. Cavalli, and C. Vaillant, Modeling epigenome folding: Formation and dynamics of topologically associated chromatin domains, Nucleic Acids Research 42, 9553 (2014).

[24] M. Chiang, D. Michieletto, C. A. Brackley, N. Rattanavirotkul, H. Mohammed, D. Marenduzzo, and T. Chandra, Polymer modeling predicts chromosome reorganization in senescence, Cell Reports 28, 3212 (2019).

[25] G. Shi and D. Thirumalai, From hi-c contact map to three-dimensional organization of interphase human chromosomes, Physical Review X 11, 10.1103/PhysRevX.11.011051 (2021).

[26] Q. MacPherson, B. Beltran, and A. J. Spakowitz, Bottom–up modeling of chromatin segregation due to epigenetic modifications, Proceedings of the National Academy of Sciences of the United States of America 115, 12739 (2018).

[27] S. Fujishiro and M. Sasai, Generation of dynamic three-dimensional genome structure through phase separation of chromatin, Proceedings of the National Academy of Sciences of the United States of America 119, e2109838119 (2022).

[28] I. Bronstein, Y. Israel, E. Kepten, S. Mai, Y. Shav-Tal, E. Barkai, and Y. Garini, Transient anomalous diffusion of telomeres in the nucleus of mammalian cells 10.1103/PhysRevLett.103.018102 (2009).

[29] H. Hajjoul, J. Mathon, H. Ranchon, I. Goiffon, J. Mozziconacci, B. Albert, P. Carrivain, J. M. Victor, O. Gadal, K. Bystricky, and A. Bancaud, High-throughput chromatin motion tracking in living yeast reveals the flexibility of the fiber throughout the genome, Genome Research 23, 1829 (2013).

[30] A. Zidovska, D. A. Weitz, and T. J. Mitchison, Micron-scale coherence in interphase chromatin dynamics., Proceedings of the National Academy of Sciences of the United States of America 110, 15555 (2013).

[31] S. C. Weber, A. J. Spakowitz, and J. A. Theriot, Nonthermal atp-dependent fluctuations contribute to the in vivo motion of chromosomal loci, Proceedings of the National Academy of Sciences of the United States of America 109, 7338 (2012).

[32] B. Gu, T. Swigut, A. Spencley, M. R. Bauer, M. Chung, T. Meyer, and J. Wysocka, Transcription-coupled changes in nuclear mobility of mammalian cisregulatory elements, Science 359, 1050 (2018).

[33] S. S. Ashwin, T. Nozaki, K. Maeshima, and M. Sasai, Organization of fast and slow chromatin revealed by single-nucleosome dynamics, Proceedings of the National Academy of Sciences of the United States of America 116, 19939 (2019).

[34] H. Ma, L.-C. Tu, Y.-C. Chung, A. Naseri, D. Grunwald, S. Zhang, and T. Pederson, Cell cycle-and genomic distance-dependent dynamics of a discrete chro-mosomal region 10.1083/jcb.201807162 (2019).

[35] H. A. Shaban, R. Barth, L. Recoules, and K. Bystricky, Hi-d: Nanoscale mapping of nuclear dynamics in single living cells, Genome Biology 21, 1 (2020).

[36] J. Lerner, P. A. Gomez-Garcia, R. L. McCarthy, Z. Liu, M. Lakadamyali, and K. S. Zaret, Two-parameter mobility assessments discriminate diverse regulatory factor behaviors in chromatin, Molecular Cell 79, 677 (2020).

[37] H. Salari, M. D. Stefano, and D. Jost, Spatial organization of chromosomes leads to heterogeneous chromatin motion and drives the liquid- or gel-like dynamical behavior of chromatin, Genome research 32, 28 (2022).

[38] A. Javer, N. J. Kuwada, Z. Long, V. G. Benza, K. D. Dorfman, P. A. Wiggins, P. Cicuta, and M. C. Lagomarsino, Persistent super-diffusive motion of escherichia coli chromosomal loci 10.1038/ncomms4854 (2014).

[39] I. Eshghi, J. A. Eaton, and A. Zidovska, Interphase chromatin undergoes a local sol-gel transition upon cell differentiation 10.1103/PhysRevLett.126.228101 (2021).

[40] A. Ghosh and N. Gov, Dynamics of active semiflexible polymers., Biophys J. 107, 1065 (2014).

[41] D. Osmanovíc and Y. Rabin, Dynamics of active rouse chains, Soft Matter 13, 963 (2017).

[42] M. Foglino, E. Locatelli, C. A. Brackley, D. Michieletto, C. N. Likos, and D. Marenduzzo, Non-equilibrium effects of molecular motors on polymers, Soft Matter 15, 5995 (2019).

[43] A. Ghosh and A. J. Spakowitz, Statistical behavior of nonequilibrium and living biological systems subjected to active and thermal fluctuations, Physical Review E 105, 14415 (2022).

[44] S. Singh and R. Granek, Active fractal networks with stochastic force monopoles and force dipoles unravel subdiffusion of chromosomal loci (2024), arXiv:2307.12310 [id=‘cond-mat.soft’ fullname =′ SoftCondensedMatter′isactive = Truealtname = Noneinarchive =′ cond mat′isg eneral = Falsedescription =′ Membranes, polymers, liquidcrystals, glasses, colloids, granularmatter′].

[45] N. Ganai, S. Sengupta, and G. Menon, Chromosome positioning from activity-based segregation, Nucl. Acids Res. 42, 4145 (2014).

[46] J. Smrek and K. Kremer, Small activity differences drive phase separation in active-passive polymer mixtures, Phys. Rev. Lett. 118, 098002 (2017).

[47] I. Chubak, S. Pachong, K. Kremer, C. Likos, and J. Smrek, Active topological glass confined within a spherical cavity, Macromolecules 55, 10.1021/acs.macromol.1c02471 (2022).

[48] A. Goychuk, D. Kannan, A. K. Chakraborty, and M. Kardar, Polymer folding through active processes recreates features of genome organization, Proceedings of the National Academy of Sciences 120, e2221726120 (2023), 10.1073/pnas.2221726120.

[49] S. Shin, G. Shi, H. W. Cho, and D. Thirumalai, Transcription-induced active forces suppress chromatin motion, Proceedings of the National Academy of Sciences 121, e2307309121 (2024), 10.1073/pnas.2307309121.

[50] J. Gelles and R. Landick, Rna polymerase as a molecular motor, Cell 93, 13 (1998).

[51] J. Ma, L. Bai, and M. D. Wang, Transcription under torsion, Science (New York, N.Y.) 340, 1580 (2013).

[52] F. R. Neumann, V. Dion, L. R. Gehlen, M. Tsai-Pflugfelder, R. Schmid, A. Taddei, and S. M. Gasser, Targeted ino80 enhances subnuclear chromatin movement and ectopic homologous recombination, Genes and Development 26, 369 (2012).

[53] J. M. Kim, P. Visanpattanasin, V. Jou, S. Liu, X. Tang, Q. Zheng, K. Y. Li, J. Snedeker, L. D. Lavis, T. Lionnet, and C. Wu, Single-molecule imaging of chromatin remodelers reveals role of atpase in promoting fast kinetics of target search and dissociation from chromatin, eLife 10, 10.7554/ELIFE.69387 (2021).

[54] M. Ganji, I. A. Shaltiel, S. Bisht, E. Kim, A. Kalichava, C. H. Haering, and C. Dekker, Real-time imaging of dna loop extrusion by condensin, Science 360, 102 (2018).

[55] S. Golfier, T. Quail, H. Kimura, and J. Brugués, Cohesin and condensin extrude dna loops in a cell-cycle dependent manner, eLife 9, 1 (2020).

[56] M. E. Cates and J. Tailleur, Motility-induced phase separation, Annual Review of Condensed Matter Physics 6, 219 (2015).

[57] D. Martin, J. O’Byrne, M. E. Cates, Étienne Fodor, C. Nardini, J. Tailleur, and F. V. Wijland, Statistical mechanics of active ornstein-uhlenbeck particles, Physical Review E 103, 10.1103/PhysRevE.103.032607 (2021).

[58] S. Brahmachari, V. G. Contessoto, M. D. Pierro, and J. D. N. Onuchic, Shaping the genome via lengthwise compaction, phase separation, and lamina adhesion, Nucleic Acids Research 50, 4258 (2022).

[59] S. Leidescher, J. Ribisel, S. Ullrich, Y. Feodorova, E. Hildebrand, A. Galitsyna, S. Bultmann, S. Link, K. Thanisch, C. Mulholland, J. Dekker, H. Leonhardt, L. Mirny, and I. Solovei, Spatial organization of transcribed eukaryotic genes, Nature Cell Biology 2022 24:3 24, 327 (2022).

[60] G. Bajpai, D. Amiad Pavlov, D. Lorber, T. Volk, and S. Safran, Mesoscale phase separation of chromatin in the nucleus, eLife 10, e63976 (2021).

[61] A. B. O. Junior, V. G. Contessoto, M. F. Mello, and J. N. Onuchic, A scalable computational approach for simulating complexes of multiple chromosomes, Journal of molecular biology 433, 10.1016/J.JMB.2020.10.034 (2021).

[62] A. W. Lau, B. D. Hoffman, A. Davies, J. C. Crocker, and T. C. Lubensky, Microrheology, stress fluctuations, and active behavior of living cells, Physical Review Letters 91, 198101 (2003).

[63] Y. Hatwalne, S. Ramaswamy, M. Rao, and R. A. Simha, Rheology of active-particle suspensions, The American Physical Society 92, 118101 (2004).

[64] F. C. Mackintosh and A. J. Levine, Nonequilibrium mechanics and dynamics of motor-activated gels 10.1103/PhysRevLett.100.018104 (2008).

[65] M. C. Marchetti, J. F. Joanny, S. Ramaswamy, T. B. Liverpool, J. Prost, M. Rao, and R. A. Simha, Hydrodynamics of soft active matter 10.1103/RevModPhys.85.1143 (2013).

[66] R. Bruinsma, A. Y. Grosberg, Y. Rabin, and A. Zidovska, Chromatin hydrodynamics, Biophysical Journal 106, 1871 (2014).

[67] A. Caspi, R. Granek, and M. Elbaum, Enhanced diffusion in active intracellular transport, Physical Review Letters 85, 5655 (2000).

[68] T. Markovich, E. Tjhung, and M. E. Cates, Chiral active matter: microscopic ‘torque dipoles’ have more than one hydrodynamic description, (2019).

[69] N. Fakhri, A. D. Wessel, C. Willms, M. Pasquali, D. R. Klopfenstein, F. C. MacKintosh, and C. F. Schmidt, High-resolution mapping of intracellular fluctuations using carbon nanotubes, Science 344, 1031 (2014).

[70] M. Guo, A. J. Ehrlicher, M. H. Jensen, M. Renz, J. R. Moore, R. D. Goldman, J. Lippincott-Schwartz, F. C. Mackintosh, and D. A. Weitz, Probing the stochastic, motor-driven properties of the cytoplasm using force spectrum microscopy, Cell 158, 822 (2014).

[71] C. Maggi, M. Paoluzzi, N. Pellicciotta, A. Lepore, L. Angelani, and R. D. Leonardo, Generalized energy equipartition in harmonic oscillators driven by active baths 10.1103/PhysRevLett.113.238303 (2014).

[72] A. Ghosh and A. J. Spakowitz, Active and thermal fluctuations in multi-scale polymer structure and dynamics, Soft Matter 18, 6629 (2022).

[73] S. Brahmachari and J. F. Marko, Chromosome disentanglement driven via optimal compaction of loop-extruded brush structures, Proceedings of the National Academy of Sciences, 201906355 (2019).

[74] J. Stenhammar, D. Marenduzzo, R. J. Allen, and M. E. Cates, Phase behaviour of active brownian particles: the role of dimensionality, Soft Matter 10, 1489 (2014).

[75] R. G. Winkler and G. Gompper, The physics of active polymers and filaments, The Journal of Chemical Physics 153, 040901 (2020), 10.1063/5.0011466/14720343/0409011online.pdf.

[76] M. Doi and S. F. Edwards, The Theory of Polymer Dynamics (Clarendon Press, 1988).

[77] J. Smrek, I. Chubak, C. N. Likos, and K. Kremer, Active topological glass, Nat. Comm. 11, 10.1038/s41467-01913696-z (2020).

[78] Y. Cui and C. Bustamante, Pulling a single chromatin fiber reveals the forces that maintain its higher-order structure, Proceedings of the National Academy of Sciences of the United States of America 97, 127 (2000).

[79] H. Yin, M. D. Wang, K. Svoboda, R. Landick, S. M. Block, and J. Gelles, Transcription against an applied force, Science 270, 1653 (1995).

[80] B. T. Donovan, A. Huynh, D. A. Ball, H. P. Patel, M. G. Poirier, D. R. Larson, M. L. Ferguson, and T. L. Lenstra, Live-cell imaging reveals the interplay between transcription factors, nucleosomes, and bursting, The EMBO Journal 38, 10.15252/embj.2018100809 (2019).

[81] D. Kurotaki, K. Kikuchi, K. Cui, W. Kawase, K. Saeki, J. Fukumoto, A. Nishiyama, K. Nagamune, K. Zhao, K. Ozato, P. P. Rocha, and T. Tamura, Chromatin structure undergoes global and local reorganization during murine dendritic cell development and activation, Proceedings of the National Academy of Sciences of the United States of America 119, e2207009119 (2022).

[82] R. Vilarrasa-Blasi, P. Soler-Vila, N. Verdaguer-Dot, N. Russiñol, M. D. Stefano, V. Chapaprieta, G. Clot, I. Farabella, P. Cuscó, M. Kulis, X. Agirre, F. Prosper, R. Beekman, S. Béa, D. Colomer, H. G. Stunnenberg, I. Gut, E. Campo, M. A. Marti-Renom José, and I. Martin-Subero, Dynamics of genome architecture and chromatin function during human b cell differentiation and neoplastic transformation 10.1038/s41467-02020849-y.

[83] B. Bonev, N. M. Cohen, Q. Szabo, L. Fritsch, G. L. Papadopoulos, Y. Lubling, X. Xu, X. Lv, J. P. Hugnot, A. Tanay, and G. Cavalli, Multiscale 3d genome rewiring during mouse neural development, Cell 171, 557 (2017).

[84] D. Michieletto, M. Chiang, D. Colí, A. Papantonis, E. Orlandini, P. R. Cook, and D. Marenduzzo, Shaping epigenetic memory via genomic bookmarking, Nucleic Acids Research 46, 83 (2018).

[85] J. A. Owen, D. Osmanovic, and L. A. Mirny, Design principles of 3d epigenetic memory systems, Science 382, 10.1126/science.adg3053 (2023).

[86] Y. Guo, E. Al-Jibury, R. Garcia-Millan, K. Ntagiantas, J. W. King, A. J. Nash, N. Galjart, B. Lenhard, D. Rueckert, A. G. Fisher, G. Pruessner, and M. Merkenschlager, Chromatin jets define the properties of cohesin-driven in vivo loop extrusion, Molecular Cell 82, 3769 (2022).

[87] N. L. Mahy, P. E. Perry, and W. A. Bickmore, Gene density and transcription influence the localization of chromatin outside of chromosome territories detectable by fish, The Journal of Cell Biology 159, 753 (2002).

[88] T. Nagano, Y. Lubling, T. J. Stevens, S. Schoenfelder, E. Yaffe, W. Dean, E. D. Laue, A. Tanay, and P. Fraser, Single cell hi-c reveals cell-to-cell variability in chromosome structure, Nature 502, 59 (2013).

[89] S. Chambeyron and W. A. Bickmore, Chromatin decondensation and nuclear reorganization of the hoxb locus upon induction of transcription, Genes development 18, 1119 (2004).

[90] R. R. Williams, S. Broad, D. Sheer, and J. Ragoussis, Subchromosomal positioning of the epidermal differentiation complex (edc) in keratinocyte and lymphoblast interphase nuclei, Experimental Cell Research 272, 163 (2002).

[91] W. Winick-Ng, A. Kukalev, I. Harabula, L. Zea-Redondo, D. Szabó, M. Meijer, L. Serebreni, Y. Zhang, S. Bianco, A. M. Chiariello, I. Irastorza-Azcarate, C. J. Thieme, T. M. Sparks, S. Carvalho, L. Fiorillo, F. Musella, E. Irani, E. T. Triglia, A. A. Kolodziejczyk, A. Abentung, G. Apostolova, E. J. Paul, V. Franke, R. Kempfer, A. Akalin, S. A. Teichmann, G. Dechant, M. A. Ungless, M. Nicodemi, L. Welch, G. Castelo-Branco, and A. Pombo, Cell-type specialization is encoded by specific chromatin topologies, Nature 2021 599:7886 599, 684 (2021).

[92] A. Agrawal, N. Ganai, S. Sengupta, and G. I. Menon, Nonequilibrium biophysical processes influence the large-scale architecture of the cell nucleus, Biophysical Journal 118, 2229 (2020).

[93] F. C. MacKintosh, Active diffusion: The erratic dance of chromosomal loci, Proceedings of the National Academy of Sciences of the United States of America 109, 7138 (2012).

[94] K. Smith, B. Griffin, H. Byrd, F. C. Mackintosh, and M. L. Kilfoil, Nonthermal fluctuations of the mitotic spindle, Soft Matter 11, 4396 (2015).

[95] T. Markovich, E. Tjhung, and M. E. Cates, Shear-induced first-order transition in polar liquid crystals, Phys. Rev. Lett. 122, 088004 (2019).

[96] D. Saintillan, M. J. Shelley, and A. Zidovska, Exten-sile motor activity drives coherent motions in a model of interphase chromatin., Proceedings of the National Academy of Sciences of the United States of America 115, 11442 (2018).

[97] P. Bera, A. Wasim, and J. Mondal, Hi-c embedded polymer model of escherichia coli reveals the origin of heterogeneous subdiffusion in chromosomal loci, Physical Review E 105, 064402 (2022).

[98] S. Chaki, L. Theeyancheri, and R. Chakrabarti, A polymer chain with dipolar active forces in connection to spatial organization of chromatin, Soft Matter 19, 1348 (2023).

[99] A. Mahajan, W. Yan, A. Zidovska, D. Saintillan, and M. J. Shelley, Euchromatin activity enhances segregation and compaction of heterochromatin in the cell nucleus, Phys. Rev. X 12, 041033 (2022).

[100] Z. Jiang, Y. Qi, K. Kamat, and B. Zhang, Phase separation and correlated motions in motorized genome, Journal of Physical Chemistry B 126, 5619 (2022).

[101] P. Eastman, J. Swails, J. D. Chodera, R. T. McGibbon, Y. Zhao, K. A. Beauchamp, L.-P. Wang, A. C. Simmonett, M. P. Harrigan, C. D. Stern, R. P. Wiewiora, B. R. Brooks, and V. S. Pande, Openmm 7: Rapid development of high performance algorithms for molecular dynamics, PLOS Computational Biology 13, 1 (2017).

